# No evidence for a bovine mastitis *Escherichia coli* pathotype

**DOI:** 10.1101/096479

**Authors:** Andreas Leimbach, Anja Poehlein, John Vollmers, Dennis Göerlich, Rolf Daniel, Ulrich Dobrindt

**Author notes:** corresponding authors Andreas Leimbach, Prof. Dr. Ulrich Dobrindt, Institute of Hygiene, Mendelstrasse 7, 48149 Münster, Germany.

## Abstract

**Background:** *Escherichia coli* bovine mastitis is a disease of significant economic importance in the dairy industry. Molecular characterization of mastitis-associated *E. coli* (MAEC) did not result in the identification of common traits. Nevertheless, a mammary pathogenic *E. coli* (MPEC) pathotype has been proposed suggesting virulence traits that differentiate MAEC from commensal *E. coli*. The present study was designed to investigate the MPEC pathotype hypothesis by comparing the genomes of MAEC and commensal bovine *E. coli*.

**Results:** We sequenced the genomes of eight *E. coli* isolated from bovine mastitis cases and six fecal commensal isolates from udder-healthy cows. We analyzed the phylogenetic history of bovine *E. coli* genomes by supplementing this strain panel with eleven bovine-associated *E. coli* from public databases. The majority of the isolates originate from phylogroups A and B1, but neither MAEC nor commensal strains could be unambiguously distinguished by phylogenetic lineage. The gene content of both MAEC and commensal strains is highly diverse and dominated by their phylogenetic background. Although individual strains carry some typical *E. coli* virulence-associated genes, no traits important for pathogenicity could be specifically attributed to MAEC. Instead, both commensal strains and MAEC have very few gene families enriched in either pathotype. Only the aerobactin siderophore gene cluster was enriched in commensal *E. coli* within our strain panel.

**Conclusions:** This is the first characterization of a phylogenetically diverse strain panel including several MAEC and commensal isolates. With our comparative genomics approach we could not confirm previous studies that argue for a positive selection of specific traits enabling MAEC to elicit bovine mastitis. Instead, MAEC are facultative and opportunistic pathogens recruited from the highly diverse bovine gastrointestinal microbiota. Virulence-associated genes implicated in mastitis are a by-product of commensalism with the primary function to enhance fitness in the bovine gastrointestinal tract. Therefore, we put the definition of the MPEC pathotype into question and suggest to designate corresponding isolates as MAEC.

## Background

Bovine mastitis is a common disease in dairy cows with a global economic impact [1]. Mastitis is an inflammation of the cow udder mostly triggered by the invasion of pathogenic bacteria, leading to reduced milk production and quality. *Escherichia coli* is a major causative agent involved in acute bovine mastitis with a usually fast recovery rate. However, in extreme cases *E. coli* mastitis can lead to severe systemic clinical symptoms like sepsis concurrent with fever [2, 3]. Occasionally, an infection with *E. coli* results in a subclinical and persistent pathology [4, 5]. Traditionally, *E. coli* associated with intramammary infections are considered to be environmental opportunistic pathogens [6]. Thus, the outcome and severity of *E. coli* mastitis was mainly attributed to environmental factors and the innate immune response of the cow reacting to pathogen-associated molecular patterns (PAMPs) (most prominently lipopolysaccharide, LPS) rather than the virulence potential of the invading strain [7]. Intramammary infusion of purified LPS induces udder inflammation symptoms similar, yet not identical, to *E. coli* invasion [7, 8]. The bovine gastrointestinal tract is a natural reservoir for commensal and pathogenic *E. coli* of high phylogenetic and genotypic diversity with the putative ability to cause mastitis [9]. Nevertheless, it was proposed that various genotypes of *E. coli* with specific phenotypes are better suited to elicit mastitis than others [10, 11, 3].

*E. coli* is a highly diverse species with commensal as well as pathogenic strains, which can colonize and persist in humans, animals, as well as abiotic environments [12, 13]. The population history of *E. coli* is largely clonal and can be structured into six major phylogenetic groups: A, B1, B2, D1, D2, and E [14, 12, 15], some publications also designate phylogroups D1 and D2 as D and F, respectively. These phylogroups have a different prevalence in various human and animal populations, but no host-restricted strains could be identified [12]. Pathogenic *E. coli* isolates are classified in different pathotypes according to the site of infection, clinical manifestation of the disease, and virulence factor (VF) repertoire. The group of intestinal pathogenic *E. coli* (IPEC) includes diarrheagenic pathotypes, which are obligate pathogens. The most prominent extraintestinal pathogenic *E. coli* (ExPEC) pathotypes are uropathogenic *E. coli* (UPEC), newborn meningitis-associated *E. coli* (MNEC), and avian pathogenic *E. coli* (APEC) [16-18]. In contrast to IPEC, which are traditionally considered to have a conserved VF repertoire, ExPEC are derived from different phylogenetic lineages and have variable VF content. Various combinations of VFs can lead to the same extraintestinal disease outcome, which solely defines an ExPEC pathotype [16, 18, 15]. However, many of these virulence-associated factors are also present in commensal strains and can be considered fitness factors (FFs), that enable or facilitate initial colonization and the establishment of an infection. These FFs have primarily evolved for gastrointestinal colonization as well as persistence, and the ability to cause extraintestinal disease is a coincidental by-product of commensalism. As a consequence, ExPEC are facultative pathogens that are recruited from the normal intestinal microbiota [19, 18, 12].

The broad spectrum of *E. coli* lifestyles and phenotypes is a result of the underlying genomic plasticity of *E. coli* strains [18]. Only up to 60% of each genome is shared by all isolates, the so-called core genome [20]. The remaining flexible genome is highly variable in individual strains. It includes genes for specific habitat adaptations or environmental conditions, and is the basis for the phenotypic diversity of *E. coli [18, 15]*. The flexible genome consists largely of mobile genetic elements (MGEs), including plasmids, genomic islands (GI), and phages, which facilitate horizontal gene transfer (HGT) and are the driving forces for microbial diversity, evolution, and adaptation potential [21].

Despite the proposal of a mammary pathogenic *E. coli* (MPEC) pathotype [3] and extensive research, no common genetic traits or VFs have been identified for *E. coli* mastitis isolates, so far [11, 22-24]. Recently, several publications analyzed *E. coli* genomes from intramammary infections, thereby expanding the method spectrum by comparative genomics approaches [25-28]. All of these studies identified various MPEC genome regions and genes with different specificity criteria and significance, many of which are not considered to be classical VFs (or even encode for unknown hypothetical functions), but also genes coding for a type VI secretion system (T6SS), LPS biosynthesis, biofilm association, metabolic functions, and the ferric iron(III)-dicitrate (Fec) uptake system. However, the studies could mostly not agree upon a common set of putative VFs, except for the Fec siderophore system. Also, these studies suffer from small genome sample size constraints, lack of phylogenetic diversity, and/or did not include commensal bovine *E. coli* comparator strains of suitable phylogenetic and genotypic diversity. So far, depending on the study, no or only one bovine commensal *E. coli* isolate has been included in these corresponding analyses [25-28].

We wanted to advance upon the previous studies by analyzing a strain panel of phylogenetic and genomic diversity comparable to *E. coli* from the bovine habitat, especially by including fecal commensal isolates from udder-healthy cows. This enables our main goal, to characterize genetic traits which define mastitis-associated *E. coli* (MAEC) in comparison to non-pathogenic commensals, while keeping track of their phylogenetic background. Putative VFs important for bovine mastitis pathogenesis should be present in the majority of mastitis isolates, regardless of phylogroup, and mostly absent in commensals. We collected a large *E. coli* VF panel from different pathotypes for detailed candidate gene and gene composition analyses. By sequencing two MAEC genomes to closure, we made it possible to analyze MGEs and evaluate their role in HGT as well as virulence of MAEC and commensal isolates. Finally, several studies suggested that mastitis virulence might have evolved in separate *E. coli* lineages and phylogroups in parallel [10, 11, 26, 27], which might involve different virulence traits and strategies. Thus, we investigated the distribution of three putative phylogroup A MPEC-specific regions from Goldstone *et al.* [26] within the phylogroups of our strain panel for pathotype association.

## Results

### Bovine-associated *E. coli* are phylogenetically highly diverse and dominated by phylogroups A and B1

We compiled a strain panel of eight MAEC and six fecal commensal strains and supplemented it with the genomes from eleven reference strains from public databases (Table 1). The reference strains are composed of eight MAEC, two fecal commensal strains, and one milk commensal strain. Serotypes were predicted *in silico* (Table 1), but could not be determined unambiguously for several draft genomes. Nevertheless, none of the analyzed strains displayed identical serotypes (except non-typable MAEC strains 131/07 and 3234/A). Thus, a correlation between certain serotypes and MAEC was not detected.

**Table 1.**
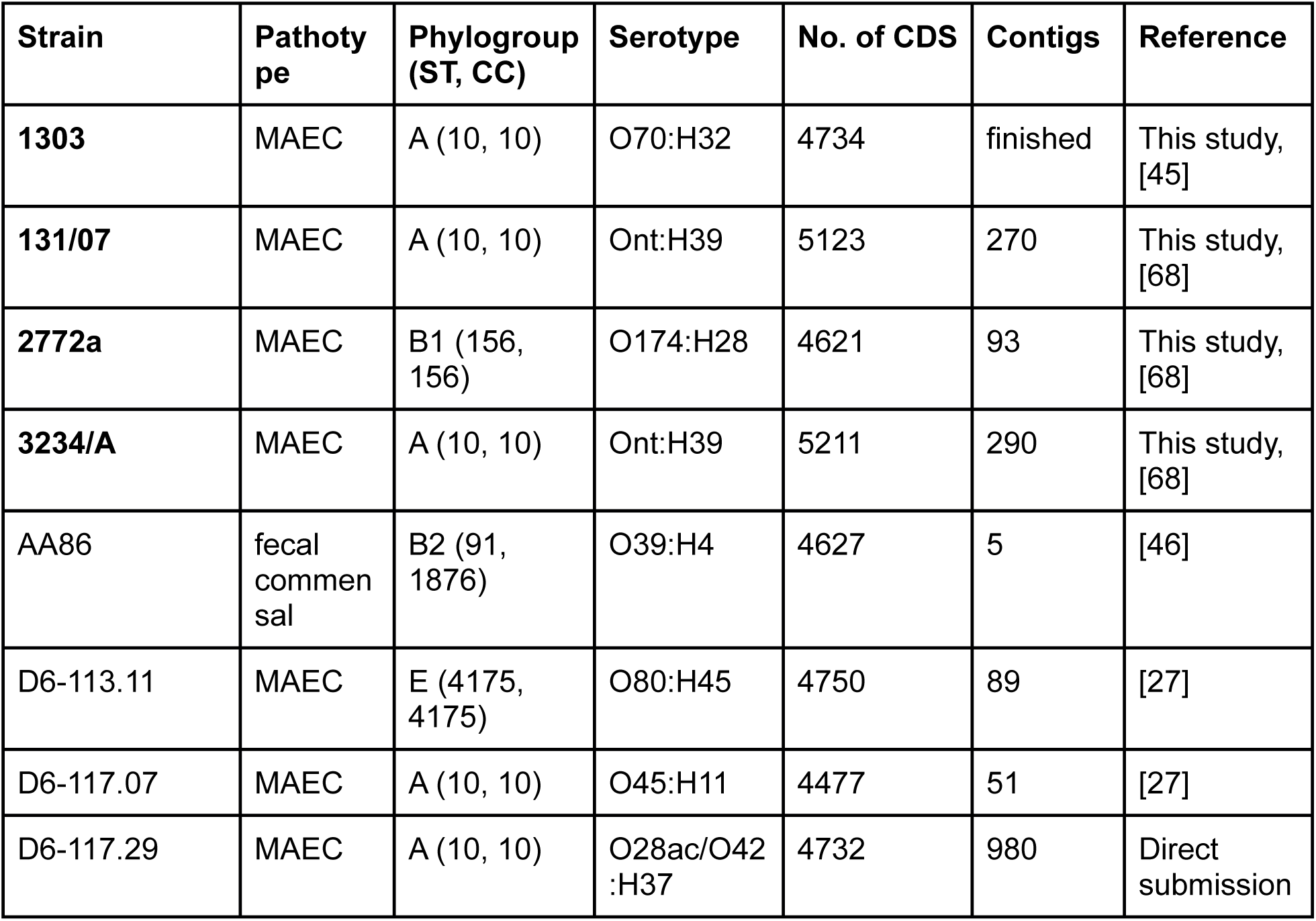
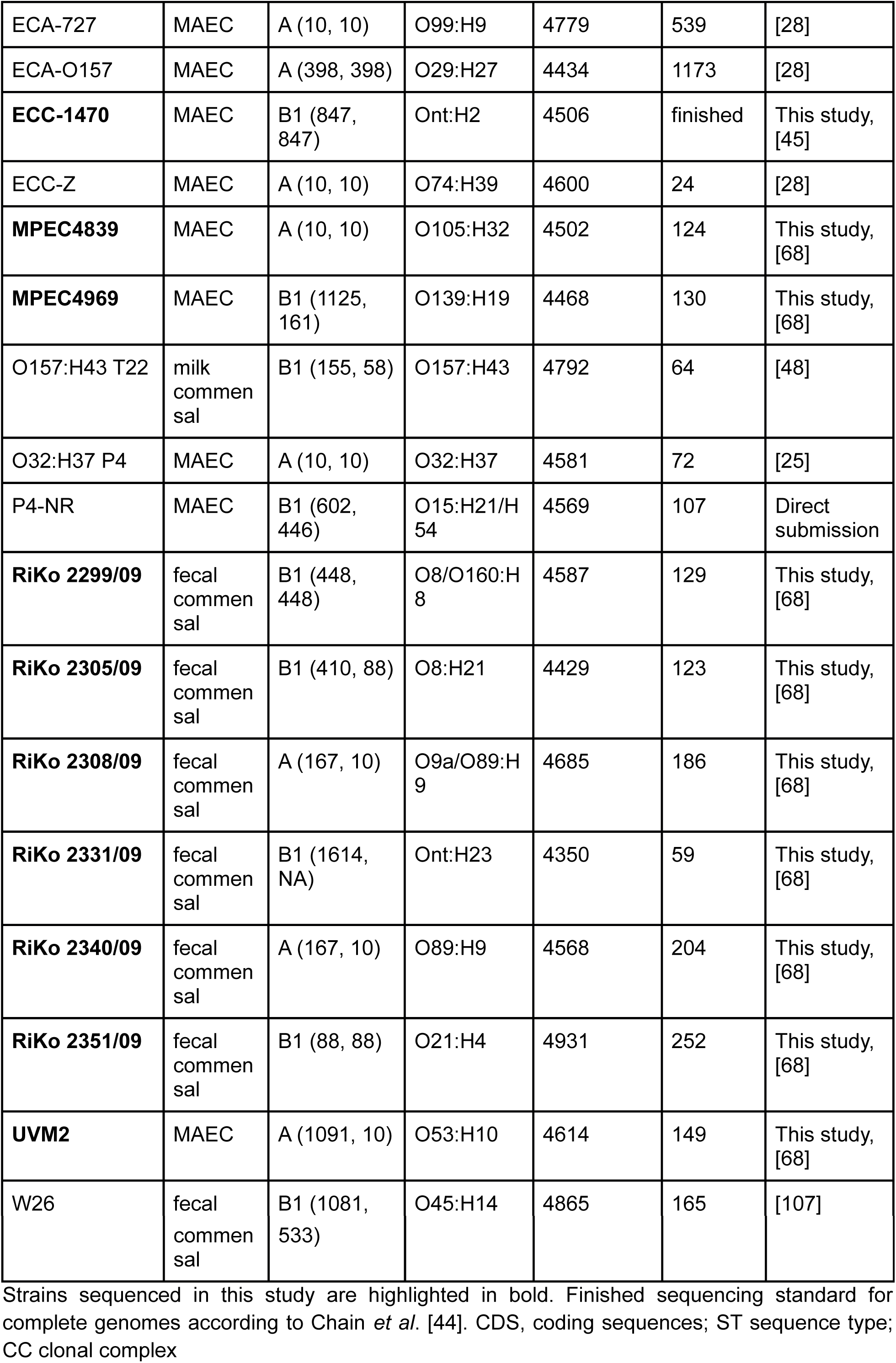
**Characteristics of the bovine-associated *E. coli* strain panel.**

The detected serotypes already indicated a high phylogenetic diversity in the strain panel. In order to obtain a more detailed view of the phylogenetic relationship of the strains, we calculated a core genome phylogeny based on a multiple whole genome nucleotide alignment (WGA) with 39 reference *E. coli* strains, four *Shigella* spp., and one *Escherichia fergusonii* strain as an outgroup. The filtered core genome WGA had a final alignment length of 2,272,130 bp, which is approximately 44% of the average *E. coli* genome size in the phylogeny (5,122,252 bp, Additional file 1 Table S1). The resulting *E. coli* population structure resolved the phylogenetic lineages described for *E. coli*, A, B1, E, D1, D2, and B2, with high bootstrap support values (Figure 1) and is in consensus with earlier studies [14, 12]. The 25 *E. coli* genomes of the bovine-associated strain panel were mostly associated with phylogroups A and B1 (13 and 10, respectively; Table 1). Most of the MAEC (11/16, 69%) belong to phylogroup A and the majority of commensal strains to group B1 (6/9, 67%). MAEC D6-113.11 and commensal AA86 are the exception to the rule by being associated with phylogroups E and B2, respectively. All phylogenetic group affiliations of the included reference strains were in accordance to their source publications (Table 1).

**Figure 1.**
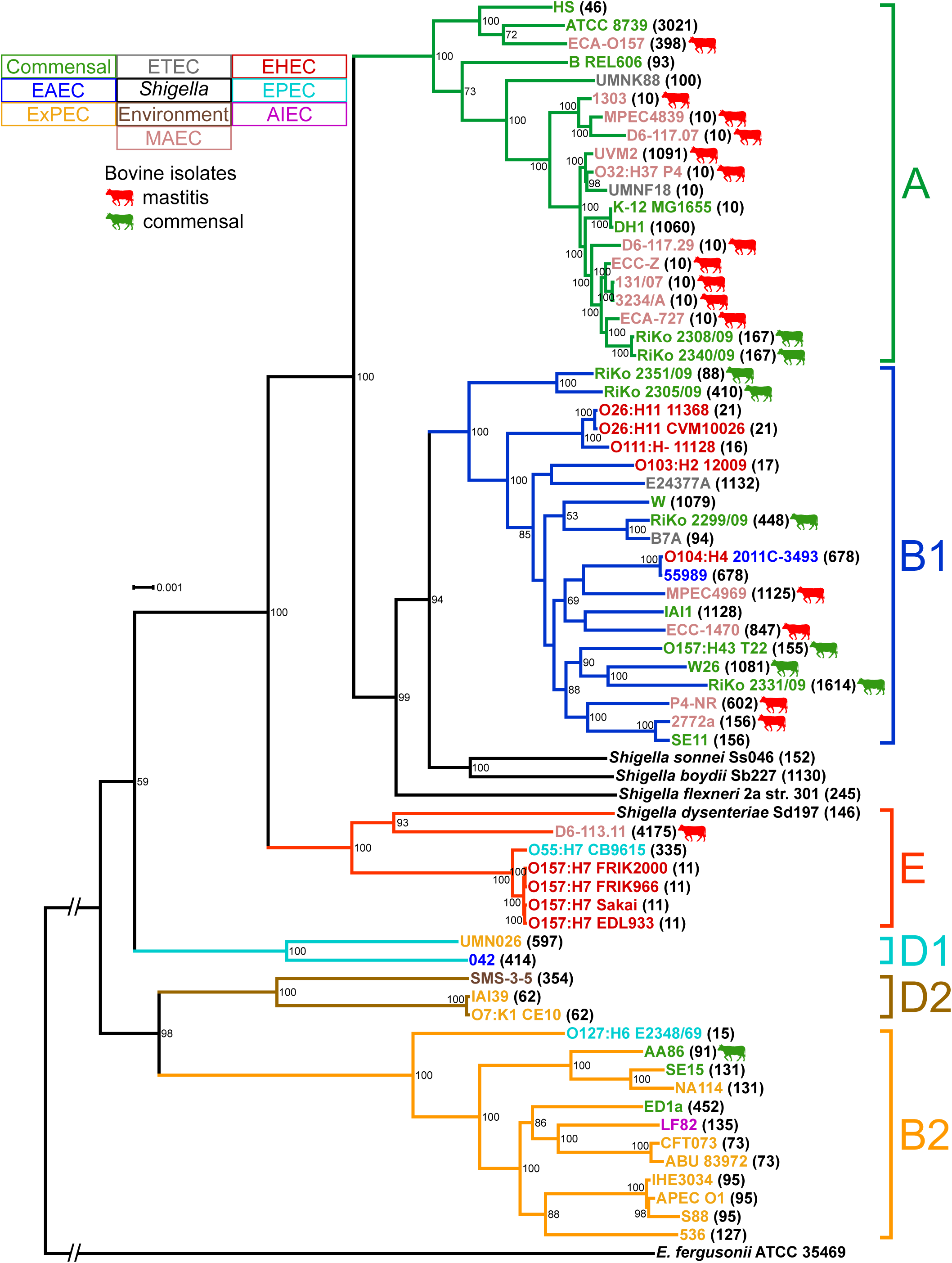
Whole genome alignment phylogeny of bovine-associated and reference *E. coli* strains. The phylogeny is based on a whole core genome alignment of 2,272,130 bp. The best scoring maximum likelihood (ML) tree was inferred with RAxML’s GTRGAMMA model with 1,000 bootstrap resamplings. The tree was visualized with Dendroscope and bootstrap values below 50 removed. *E. fergusonii* serves as an outgroup and the corresponding branch is not to scale. Bovine-associated *E. coli* are indicated by colored cows, and both *E. coli* pathotypes and phylogroups are designated with a color code. ST numbers from the MLST analysis for each strain are given in parentheses. *E. coli* isolated from cows are widely distributed in the phylogroups and both commensal and MAEC strains are interspersed in the phylogenetic groups with a polyphyletic history.

To enhance backwards compatibility, we determined the sequence types (ST) for all strains analyzed in the WGA phylogeny according to the Achtman *E. coli* multi-locus sequence typing (MLST) scheme (Additional file 2: Table S2) [13]. The calculated minimum spanning tree (MST) supports the phylogenetic history depicted in the WGA phylogram and confirms the diversity of bovine-associated *E. coli* (Additional file 3: Figure S1). ST10, with nine occurrences, is the most common ST in the 25 *E. coli* genomes from the bovine-associated strain panel. In fact, all bovine-associated *E. coli* of phylogroup A are members of clonal complex 10 (CC10), except for *E. coli* ECA-O157 (ST398, CC398). Nevertheless, the majority of the 25 *E. coli* genomes have different STs, corroborating their phylogenetic diversity.

### Gene content correlates with phylogenetic lineages of bovine-associated *E. coli*

Despite the phylogenetic diversity of the bovine-associated *E. coli*, we were interested to see if functional convergence of bovine MAEC or commensals exists. There might be a defining subset of genes or VFs for MAEC from different phylogenetic backgrounds that would point to a putative MPEC pathotype. For this purpose we determined the similarity of the genomes based on the presence/absence of all orthologous groups (OG) calculated for the strain panel. Such an analysis, visualized as a so-called gene content tree, has the advantage of considering the core as well as the flexible genome, in contrast to the WGA core genome phylogeny (in which the flexible genome is intentionally filtered out in order to maximize the robustness of the inferred phylogenetic history). Thus, this method can be used to detect functional similarities based on similar gene content. We clustered all strains based on gene content by calculating the best scoring maximum likelihood (ML) tree of the binary matrix representing the presence and absence of OGs (Additional file 4: Dataset S1). The topology of the resulting gene content tree mirrors the phylogenetic lineages of the WGA phylogeny with high analogy (Figure 2). All bifurcations that define phylogroups in the gene content tree have high bootstrap values. For comparison purposes we visualized the high similarity between WGA genealogy and gene content tree in a tanglegram (Additional file 3: Figure S2A and B). This diagram shows that not only the phylogroups are conserved, but also the phylogenetic relationships between individual *E. coli* isolates within the phylogroups. However, some minor differences in the bifurcations between phylogeny and gene content clustering were detected. The two biggest differences concern the placement of phylogroups B2/E and MAEC ECA-O157. In contrast to the WGA-based phylogeny, which clusters phylogroups B2 and E outside the A/B1 sister taxa, the gene content dendrogram places these phylogroups closer to B1 than A (Figure 2 and Additional file 3: Figure S2B). This appears to be due to a more similar gene content, as phylogroups B2/E have a higher recombination frequency with phylogroup B1 than with A [29, 30]. Strain ECA-O157 represents an outlier branch in comparison to all other included *E. coli* genomes based on gene content (Figure 2). As this strain is the only strain in phylogroup A that does not belong to the closely related CC10 cluster, this explains its gene content divergence to the other A strains in the gene content tree, which is also apparent in the WGA core genome phylogeny. However, the outlier-clustering of ECA-O157 might also be a result of the high fragmentation of the draft genome (nearly 1000 contigs > 500 bp, Additional file 5: Table S3) and the resulting uncertain accuracy of CDS (coding DNA sequence) predictions on which OG analyses are dependent.

**Figure 2.**
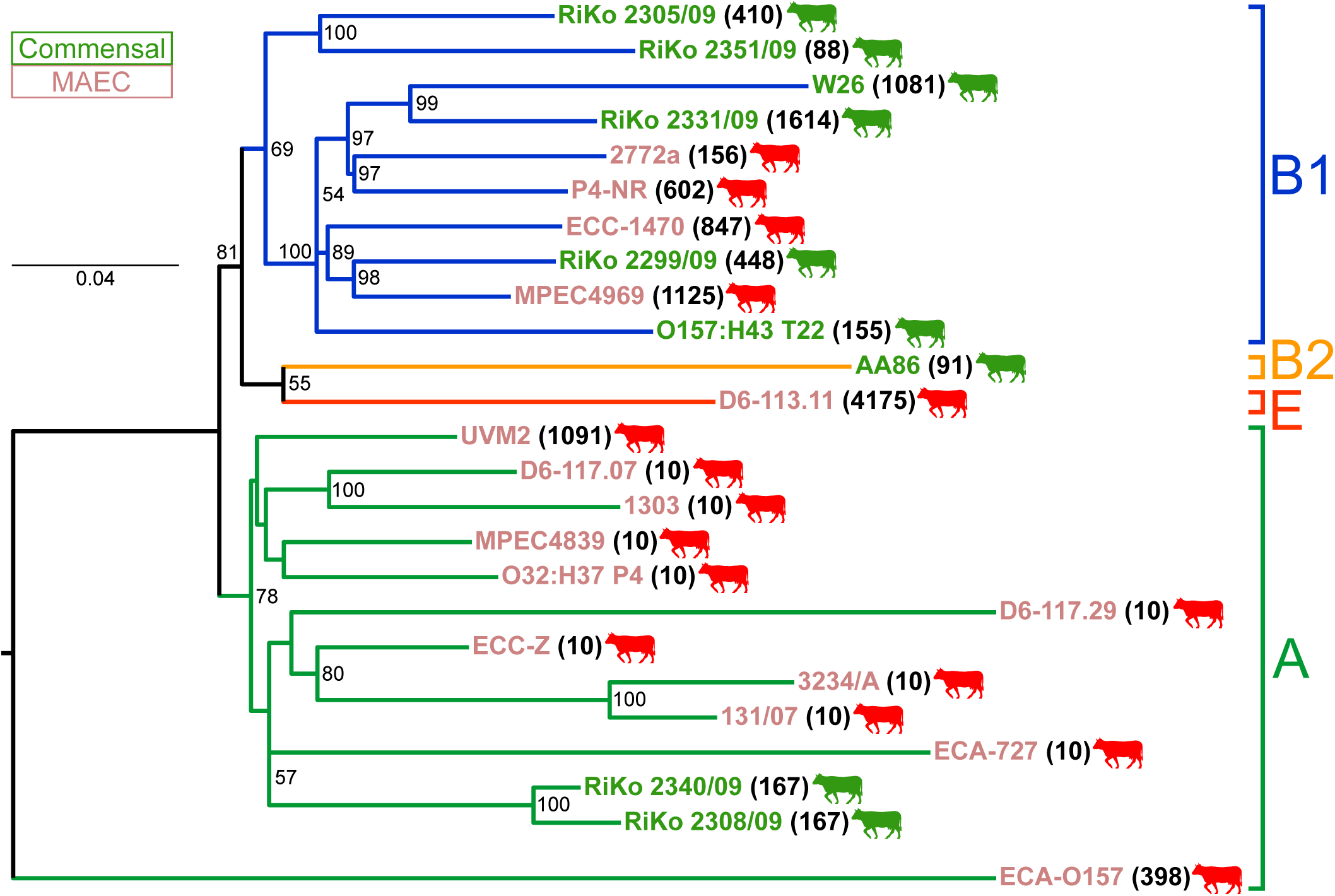
Gene content clustering tree of the bovine-associated *E. coli*. The gene content best scoring ML dendrogram is based upon the presence or absence of orthologous groups (OGs) with 1000 resamplings for bootstrap support values. The tree was visualized midpoint rooted with FigTree and bootstrap values below 50 removed. The distance between the genomes is proportional to the OGs present or absent. The tree topology of the gene content tree follows closely the core genome WGA phylogeny. There is no functional convergence between MAEC or commensal strains, rather a highly diverse gene content.

In conclusion, no functional convergence of bovine MAEC or commensals could be detected and the phylogenetic diversity of the strains is also apparent in a highly diverse gene content.

### MAEC possess no virulence-attributed orthologs in comparison to commensal strains

Since no large scale gain or loss of bovine MAEC-or commensal-associated genes could be detected in the gene content tree, we looked into the distribution of OGs in more detail, in order to search for genotypic traits enriched in bovine mastitis or commensal isolates. From our point of view, only the comparison of a larger set of MAEC genome sequences with that of bovine commensals is suitable to address this question. If any VFs/FFs existed, that play an important role in the pathogenesis of MAEC, we would expect a wide distribution of the encoding genes among MAEC strains compared to commensals.

The pan-genome of the 25 bovine-associated *E. coli* strains amounted to 116,535 CDS and a total of 13,481 OGs using BLASTP+ with 70% identity and coverage cutoffs. Because of the open nature of the *E. coli* pan-genome [31], all genomes included OGs, which were absent in every other compared strain (so-called singletons; Additional file 6: Dataset S2). The largest numbers of singletons were detected in the highly fragmented genomes of strains D6-117.29 (n=455), ECA-O157 (n=865), and ECA-727 (n=615), a likely consequence of the high number of contig ends and uncertain open reading frame (ORF) predictions (Additional file 5: Table S3). Also, large numbers of singletons in genomes AA86 (n=422) and D6-113.11 (n=361) are to be expected, as these are the only compared genomes of their respective phylogroups, B2 and E. The majority of singletons encode typical proteins of the flexible genome, like hypothetical proteins, proteins associated with MGEs (transposases, phages), restriction modification systems, O-antigen biosynthesis, CRISPR, conjugal transfer systems, and sugar transport/utilization operons. Although several of these genes and gene functions have previously been identified as MAEC-associated in small strain panels [25, 27], they most likely play no role in mastitis because of their presence in commensals and/or low prevalence in MAEC.

To determine OGs which are characteristic of mastitis-associated or bovine commensal isolates, we screened the 13,481 OGs of the bovine-associated *E. coli* pan-genome for OGs which are significantly (p < 0.05) associated with one of these two groups of strains, using Fisher’s exact test. 240 OGs displayed a significant association with a pathotype. However, none of these OGs remained significantly associated with either mastitis or commensal isolates when a Bonferroni correction was applied (Figure 3A, Additional file 3: Figure S3A). Furthermore, none of these OGs were exclusively present in all mastitis, but absent from all commensal isolates tested and vice versa. In order to identify OGs with a wide distribution in one pathotype in comparison to the other, we looked for OGs present in at least 70% of the genomes of one pathotype and maximally in 30% of the other. This resulted in 36 “MAEC-” and 48 “commensal-enriched” OGs, most of which displayed a significant association (Figure 3A and Additional file 7: Dataset S3).

**Figure 3.**
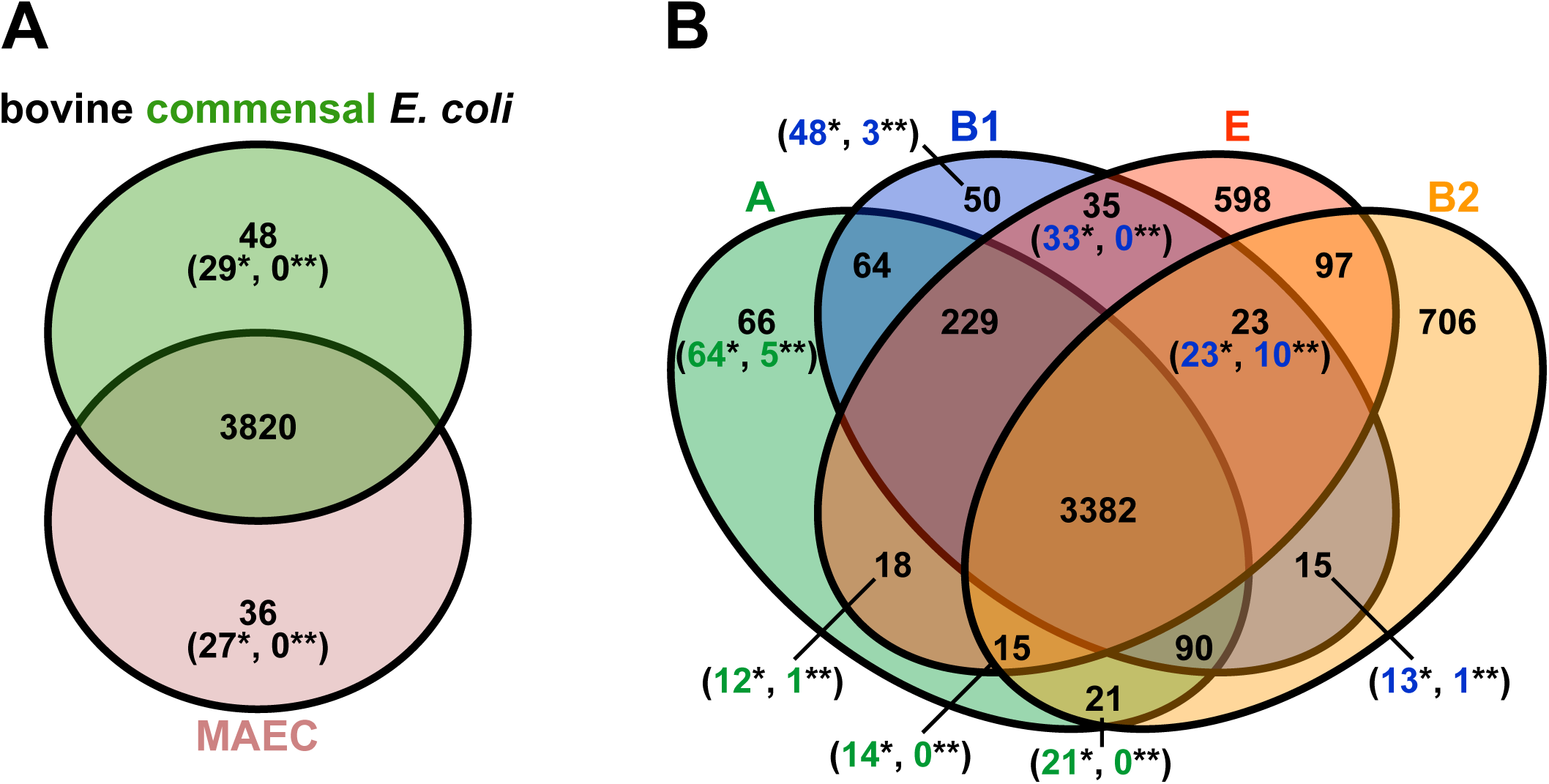
Venn diagrams for gene family enrichment in pathotypes or phylogroups. A) Enrichment of OGs in pathotypes (MAEC or commensal) was determined statistically (numbers in parentheses; Fisher’s exact test, p < 0.05) after applying 70% inclusion and 30% exclusion group cutoffs (numbers without parentheses). Numbers with a single asterisk correspond to OGs with a statistically significant association while two asterisks indicate remaining significant associations after a Bonferroni correction. B) Enrichment of OGs in phylogenetic groups (A, B1, B2, or E) was determined based on 70% inclusion and 30% exclusion group cutoffs. Statistic testing for OG association (Fisher’s exact test, p < 0.05) was performed only for the multi-genome phylogroups A versus B1. Only very few OGs could be detected as pathotype enriched. Instead, OG distribution is strongly affected by phylogenetic background.

Because phylogeny was shown to exhibit a strong effect on the gene content of *E. coli* isolates and shared ancestry might overshadow functional relatedness, we tested the 13,481 OGs additionally for a significant association with the phylogroups A or B1. This resulted in 410 significantly associated OGs. After Bonferroni correction, six OGs remained significantly associated to phylogroup A, whereas 14 OGs remained significantly phylogroup B1-associated (Additional file 3: Figure S3B). We used the same inclusion and exclusion cutoffs to identify Ogs that were enriched in genomes of the four phylogroups (A, B1, B2, and E; Figure 3B and Additional file 8: Dataset S4). This analysis resulted in many phylogroup-enriched OGs, supporting the impact of phylogeny on gene content and the similarity between the gene content tree and WGA phylogeny. An “all-strain” soft core genome (as defined by Kaas *et al.* [20]) with this 70% inclusion cutoff included 3,842 OGs, which is about 82% of the average number of CDS in the genomes (Additional file 9: Table S4).

#### Commensal-enriched orthologous groups are associated with fitness factors

Of the 29 significant commensal-enriched OGs, eight OGs are not simultaneously enriched in a phylogroup (Additional file 7: Dataset S3). These include the aerobactin siderophore biosynthesis operon (*iucABCD*) with the siderophore receptor-encoding (*iutA*) and the associated putative transport protein ShiF-encoding genes (locus tags in strain RiKo 2340/09 RIKO2340_186c00010 to RIKO2340_186c00060) as well as two OGs coding for IS element (insertion sequence)-related proteins (paralogs RIKO2340_128c00050/RIKO2340_203c00010 and RIKO2340_203c00020). 20 of the 21 remaining commensal-enriched OGs are significantly associated with phylogroup B1 and include fimbrial genes, genes of the galactitol (*gatZCR*) phosphotransferase systems (PTS) as well as genes for sucrose catabolism (*cscKA*), a putative ABC transporter, the ChpB-ChpS toxin-antitoxin system, and a lipoprotein. Their role for bovine commensalism remains unclear, especially because of their additional association with phylogroup B1.

#### MAEC-enriched orthologous groups are mostly associated with mobile genetic elements

Thirteen of the 27 significantly mastitis-associated OGs are significantly enriched in phylogroup A, two are present in the soft core genome, and two are absent in the phylogroup B2 (commensal AA86). The remaining ten mastitis-associated OGs, which do not display phylogenetic or core enrichment, do not include any coherent gene cluster (Additional file 7: Dataset S3). However, eight of them are located in close proximity to each other in the genome of strain 1303 (*rzpQ* EC1303_c16730, *ydfR* EC1303_c16750, *quuQ_1* EC1303_c16790 (paralog to *quuQ*_2 in 1303 prophage 4), *relE* EC1303_c16830, *relB* EC1303_c16840, *flxA* EC1303_c16860, putative integrase EC1303_c16890, and hypothetical protein EC1303_c16900). All of these proteins belong to a prophage without noticeable features (see *E. coli* 1303 prophage 2 below). Additionally, three genes included in the soft core and enriched in phylogroups A/B1/E, *cspI*, *cspB* (“cold shock proteins”, EC1303_c16710 and EC1303_c16770, respectively), and *recE* (“exonuclease VIII, 5’ -> 3’ specific dsDNA exonuclease“, EC1303_c16970) also lie within the same prophage region. Because the *E. coli* 1303 prophage 2 genome does not contain genes related to metabolic or virulence functions, the role of the respective encoded gene products in mastitis cannot be determined. The last two OGs without phylogroup or core enrichment, *ylbG* (E1470_c05180) and *ybbC* (EC1303_c04920), encode for a putative DNA-binding transcriptional regulator and a putative immunity protein, respectively, and are associated with an *rhs* element. The 13 OGs that are also significantly enriched in phylogroup A encode for a transcriptional regulator (*rmhR* EC1303_c24270), an alpha amylase (EC13107_63c00240), a toxin/antitoxin system (*yafNO*, EC1303_c02750 and EC1303_c02760), a lipoprotein (*ybfP* EC1303_c06580), a phsohpodiesterase (*yaeI* EC1303_c01600), a malonyl CoA-acyl carrier protein transacylase (*ymdE* EC1303_c10470), a transposase (*insL*1_2 gene EC1303_c28750), and hypothetical proteins. According to the sequence contexts in these strains, the genes cannot be unambiguously localized in prophage regions or typical pathogenicity islands. Additionally, *eprI* (EC1303_c29770) encodes a type III secretion apparatus inner ring protein and is associated with a pathogenicity island (PAI). Finally, two genes contained in MAEC 1303 prophage 1, encoding for an exonuclease (EC1303_c12230) and an envelope protein (EC1303_c12530), are also associated with phylogroups A/B1/E. In summary, the putative mastitis-eliciting function of any of the genes within the significantly MAEC-associated OGs is unclear. A truly meaningful correlation between OGs and pathotypes (mastitis vs. commensal) could not be observed. Instead, several OGs are significantly associated with phylogroups A or B1. No traditional *E. coli* VFs have been found among MAEC-enriched OGs.

### Genomic islands and prophages in MAEC 1303 and ECC-1470 contain only few well-known virulence-associated genes

Both finished *E. coli* 1303 and ECC-1470 genomes include several putative pathogenicity, resistance, and metabolic islands, as well as prophages (Additional file 10: Dataset S5 and Additional file 11: Dataset S6). GIs could only be detected in the chromosomes of the closed genomes, but not on the respective plasmids. However, on the F plasmid present in *E. coli* 1303, p1303_109, a smaller 17-kb transposable element was identified. Mastitis isolate 1303 additionally harbors an episomal circularized P1 bacteriophage [32], designated p1303_95. Generally, the genome of mastitis isolate 1303 includes twelve GIs ranging in size from 11 to 88 kb and encoding from 11 to 81 CDSs (Additional file 10: Dataset S5). One large composite GI (GI4) combines pathogenicity-and resistance-related genes. It partly contains the biofilm-associated polysaccharide synthesis *pga* locus. The resistance-related genes of GI4 are located on the AMR-SSuT (antimicrobial multidrug resistance to streptomycin, sulfonamide, and tetracycline) island, which is prevalent in *E. coli* from the bovine habitat [33, 34]. The encoded resistance genes are *strAB*, *sul2*, and *tetDCBR*. A comparison of the corresponding genomic region of *E. coli* 1303 with two publicly available AMR-SSuT island sequences is shown in Additional file 3: Figure S4. Transposon Tn*10*, also present on the resistance plasmid R100, is an integral part of the AMR-SSuT island and comprises the *tetDCBR* genes. This highlights the composite nature of the AMR-SSuT island and of GI4 in general. The resistance markers of AMR-SSuT are prevalent, as seven strains of the panel contain some or all of the genes (D6-117.29, ECA-727, RiKo 2305/09, RiKo 2308/09, RiKo 2340/09, RiKo 2351/09, and W26).

The twelve GIs harbored by mastitis isolate ECC-1470 vary in size between 8 to 58 kb and code for 9 to 61 CDSs (Additional file 11: Dataset S6). *E. coli* ECC-1470 (Ont:H2) encodes for a flagellin of serogroup H2 and an uncharacterized small alternative flagellin, FlkA, encoded on GI10. The neighbouring *flkB* gene encodes for a FliC repressor. This small alternative flagellin islet can elicit unilateral H-antigen phase variation [35, 36]. The MAEC strain P4-NR (O15:H21/H54), which usually expresses a serotype H21 flagellin, also harbours a similar alternative flagellin system determinant consisting of the serotype H54 flagellin gene *flmA54* and the associated *fliC* repressor-encoding gene *fljA54*. GI12 of ECC-1470 is a large PAI containing a fimbrial operon of the P adhesin family (*pixGFJDCHAB*, *pixD* is a pseudogene), a phosphoglycerate transport operon (*pgtABCP*), the putative MAEC-associated Fec transport operon (*fecEDCBARI*), the 9-O-acetyl-*N*-acetylneuraminic acid utilization operon (*nanSMC*), and the type 1 fimbriae operon (*fimBEAICDFGH*). This PAI is a composite island with the 5’-end similar to PAI V from UPEC strain 536 with the *pix* and *pgt* loci, also present in human commensal *E. coli* A0 34/86 [37, 38], and the 3’-end similar to GI12 of MAEC 1303 with the *nan* and *fim* gene clusters. *E. coli* ECC-1470 GI4 codes for a lactose/cellobiose PTS system (*bcgAHIFER*, *bcgI* is a pseudogene).

Four prophages were predicted in the genome of MAEC 1303 ranging from 29 to 48 kb encoding for 44 to 59 CDSs (Additional file 10: Dataset S5). These prophage genomes do not comprise many virulence-associated genes, and mostly code for functions required for maintenance and mobilization. The only exception is *bor*, a gene of phage lambda widely conserved in *E. coli* and encoded by strain 1303 chromosomal prophage 1. The outer membrane lipoprotein Bor is homologous to Iss (increased serum survival) and involved in serum resistance of ExPEC [39, 40]. The lack of putative *E. coli* VFs encoded by prophages is also true for the five predicted prophage genomes of MAEC ECC-1470 (Additional file 11: Dataset S6). Two outer membrane proteins (OMPs) are encoded by ECC-1470 prophage 1, the porin NmpC and the omptin OmpT.

In summary, the MGEs of MAEC strains 1303 and ECC-1470 do not carry many known virulence-associated genes, which may entail an advantage to mastitis pathogens. To illustrate the resulting mosaic-like structure of *E. coli*, we created circular genome diagrams for all MAEC 1303 and ECC-1470 replicons indicating the core and the flexible genome by labeling the predicted GIs and prophages (Additional file 3: Figure S5A and B). Importantly, the prevalence and dissemination of the MGEs were not correlated with the pathotypes.

### Virulence or fitness factors present in bovine commensal *E. coli* or MAEC

To examine the distribution of virulence-associated factors in more detail we searched for well-known *E. coli* VFs encoded by the bovine-associated *E. coli* genomes (Additional file 12: Table S5) [41]. Only about half of the 1,069 gene products involved in the biosynthesis and function of 200 *E. coli* virulence and fitness-associated factors yielded BLASTP+ hits in the 25 bovine-associated *E. coli* genomes. Virulence-associated proteins of the VF panel present in (556) and absent from (513) these *E. coli* genomes are listed in Additional file 12: Table S5. Results of the BLASTP+ hits for the virulence-associated proteins are listed in Additional file 13: Table S6. Many classical IPEC VFs [42] were not present in the bovine-associated strains. Interestingly, all major virulence factors of EHEC are missing. Furthermore, several VFs associated with ExPEC [17] were absent, such as several typical serine protease autotransporters of *Enterobacteriaceae* (SPATE) like Sat and Pic (type V secretion systems, T5SS), S fimbriae, salmochelin siderophore, colicin V, and colibactin. The fecal isolate RiKo 2351/09 of phylogroup B1 yielded the most virulence-associated protein hits (297), whereas MAEC ECA-O157 of phylogroup A the fewest (162). There were 241 virulence-associated protein hits on average in the strains included in this study. We could not detect a correlation between the number of virulence-associated genes and pathotype as both, commensal strains and MAEC, exhibited comparable average virulence-associated genes hits (250 and 237, respectively). The average number of virulence-associated genes was in the same range in the *E. coli* genomes of the different phylogroups (phylogroup A: 233, B1: 254, B2: 227, and E: 235).

We converted the BLASTP+ VF hits for each strain into a presence/absence binary matrix (Additional file 14: Dataset S7) to enable grouping of the compared strains according to their VF content (Additional file 3: Figure S2C). Most of the genomes belonging to the same phylogroup clustered together. However, the phylogenetic relationships of the strains from the gold standard WGA phylogeny are not all retained. Consequently, the association of the strains with phylogroups in the VF content tree is not as well conserved as in the overall gene content tree, as shown by a tanglegram with the WGA phylogeny (Additional file 3: Figure S2D). The presence and absence of the VFs in the different strains were visualized in a heatmap in which the respective genome columns are ordered according to the clustering results (Figure 4). This heatmap is replicated with the corresponding virulence-associated gene names in Additional file 3: Figure S6. Analogous to the all-strain soft core genome we determined an “all-strain” soft core VF set. In consideration that many fragmented draft genomes are included in the strain panel, we once more applied a 70% inclusion cutoff. As a result, virulence-associated genes were included if they were present in at least 18 of the 25 bovine *E. coli* genomes analyzed. The resulting 182 virulence-associated genes (Additional file 15: Table S7) included determinants generally considered to be widely present in *E. coli* isolates, like the Flag-1 flagella system, the operons encoding type 1 fimbriae, and the *E. coli* common pilus (ECP). But also curli fimbriae, the lipoprotein NlpI, outer membrane protein OmpA, and several iron transport systems (ferrous low pH (*efe*/*ycd*), enterobactin (*ent*, *fes*, and *fep*), ferrous (*feo*), and ferrichrome (*fhu*)) are included. Additionally, several T2SS genes, 16 of the 32 *E. coli* type three secretion system 2 (ETT2) genes, and two genes from the ECC-1470 T6SS/1, *impA* and a gene coding for a Hcp T6SS effector-like protein (E1470_c02180), are enclosed.

**Figure 4.**
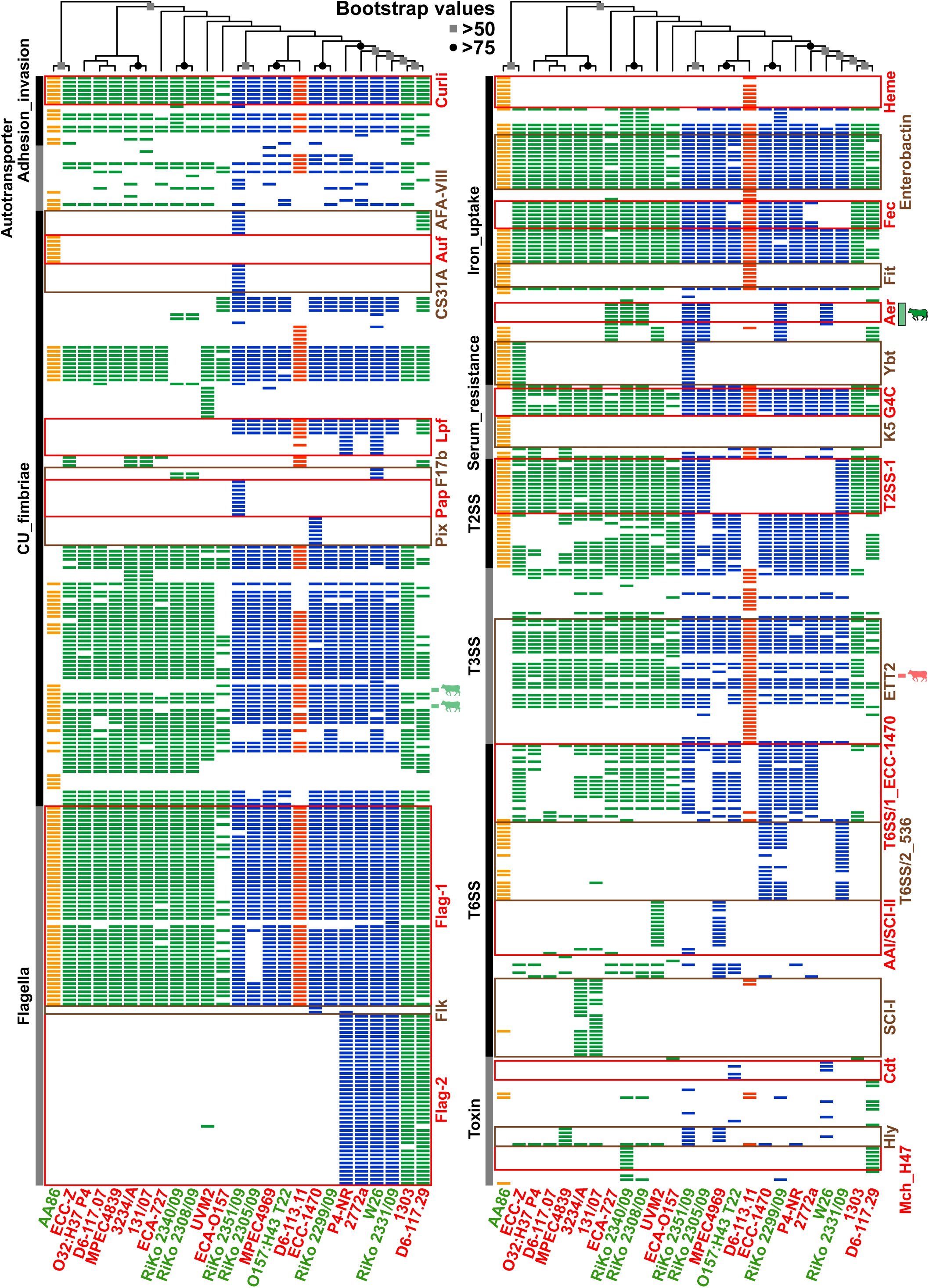
Heatmap indicating presence or absence of virulence factors. Each row of the binary matrix indicates the presence or absence of a virulence-associated gene (a BLASTP+ hit). VF classes are indicated at the side in black and grey. Strain names are color-coded for MAEC (red) or commensal (green) pathotype affiliation, columns for strain phylogroup affiliation (green: A, blue: B1, orange: B2, red: E). The clustering dendrogram attached to the heatmaps is based upon the whole binary dataset (not for each heatmap separately) of a best scoring ML tree with 1000 bootstrap resamplings (a more detailed representation of the cladogram can be found in Additional File 3: Figure S2C). Bootstrap support values are arbitrarily indicated at the bifurcations of the cladogram. Statistically significant pathotype-enriched VF genes are indicated for MAEC and commensal isolates by cows in red or green, respectively. Only the aerobactin biosynthesis cluster (Aer) plus transport protein ShiF is significantly commensal-enriched and not associated with a phylogroup (indicated by a black-rimmed and opaque green cow). All other pathotype-enriched virulence-associated genes also have a significant phylogroup association. The genes of well-known and important *E. coli* VFs are highlighted in alternating red and brown squares: Curli = curli fibres, AFA-VIII = aggregative adherence fimbriae AFA-VIII, Auf = fimbrial adhesin, CS31A = CS31A capsule-like antigen (K88 family adhesin), Lpf = long polar fimbriae, F17b = F17b fimbriae, Pap = P/Pap pilus, Pix = Pix fimbriae, Flag-1 = *E. coli* peritrichous flagella 1 gene cluster, Flk = alternative *flk* flagellin islet, Flag-2 = *E. coli* lateral flagella 2 gene cluster, Heme = *chu* heme transport system, Enterobactin = enterobactin biosynthesis/transport gene cluster, Fec = ferric iron(III)-dicitrate uptake system, Fit = ferrichrome iron transport system, Aer = aerobactin biosynthesis cluster with *iutA* receptor, Ybt = yersiniabactin iron transport system, G4C = group 4 capsule, K5 = K5 capsule, T2SS-1 = *gsp* general secretion pathway 1, ETT2 = *E. coli* type three secretion system 2, T6SS/1_ECC-1470 = MAEC ECC-1470 subtype i1 T6SS/1, T6SS/2_536 = UPEC 536 subtype i2 T6SS 2, AAI/SCI-II = EAEC 042 subtype i4b T6SS 3, SCI-I = EAEC 042 subtype i2 T6SS 2, Cdt = cytolethal distending toxins, Hly = alpha-hemolysin, Mch_H47 = microcin H47. Clustering of the strains according to virulence-associated gene presence/absence also follows mostly the phylogenetic history of the strains, no clustering of pathotypes was detected. Both MAEC and commensal isolates are distinguished by the lack of classical pathogenic *E. coli* VFs. The same heatmap, but including gene names/locus tags, can be found in Additional File 3: Figure S6.

In conclusion, the VF variety observed is in accordance with the high diversity of bovine-associated *E. coli*.

### Specific virulence or fitness genes cannot be unambiguously detected for MAEC or commensal bovine isolates

According to Fisher’s exact test performed with the 556 VF-related genes detected in our strain panel, 30 were significantly associated with mastitis or commensal isolates (Additional file 16: Dataset S8). However, with a Bonferroni correction for multiple comparisons we could not detect a significant association (Additional file 3: Figure S3C), also no VF was exclusively present in MAEC or commensal isolates. Nine virulence genes were significantly associated with mastitis genomes. Although overrepresented in mastitis isolates, the *fecRIABCDE* genes as well as the type 1 fimbriae minor subunit-encoding gene *fimG* are also present in at least 50% of the commensal genomes analyzed. The only MAEC-enriched virulence-associated OG that fulfills the 70%/30% inclusion/exclusion cutoffs, was the ETT2 *eprI* gene. Nevertheless, this gene is also enriched in phylogroups A/E (Figure 5A, Table 2, and Additional file 17: Dataset S9). Additionally, *eprI* and the *fecBCDE* genes were also tested significantly enriched in phylogroup A strains in comparison to B1 strains; *eprI* even with a Bonferroni correction. Overall, 58 VF-related genes were significantly associated with phylogroup A or B1, and of these six with phylogroup A and 12 with phylogroup B1 after a Bonferroni correction (Additional file 3: Figure S3D and Additional file 17: Dataset S9). 21 virulence-associated genes were associated with commensal strains. Seven of them, including EC042_1639 and *ydeT* (coding for parts of Yde fimbriae), *gspFHI* and *yghJ* (coding for components of the T2SS-2 system), as well as the T3SS effector-encoding *espX1* gene display at the same time a significant enrichment in phylogroup B1 strains. Of these, the Yde fimbrial genes are significantly phylogroup B1-associated also with a Bonferroni correction and fulfill the 70%/30% inclusion/exclusion cutoff for phylogroups B1/B2/E (Table 2). The residual fourteen VF genes significantly associated with commensal isolates were not phylogroup-enriched. These virulence-associated genes are involved in biosynthesis and transport of the aerobactin siderophore (*iucABCD*, *iutA*, *shiF*), F17 fimbriae biogenesis (f17d-C, pVir_8, pVir_9) or code for an enterotoxin (*senB*) and colicin-related functions (*cjrABC, imm*). The presence of the aerobactin genes were also within the 70%/30% inclusion/exclusion cutoffs and not simultaneously enriched in a phylogroup (Table 2 and Additional file 16: Dataset S8). Altogether, most of the significantly phylogroup-associated VFs were also included with the 70%/30% cutoffs (Figure 5B and Additional file 17: Dataset S9).

**Figure 5.**
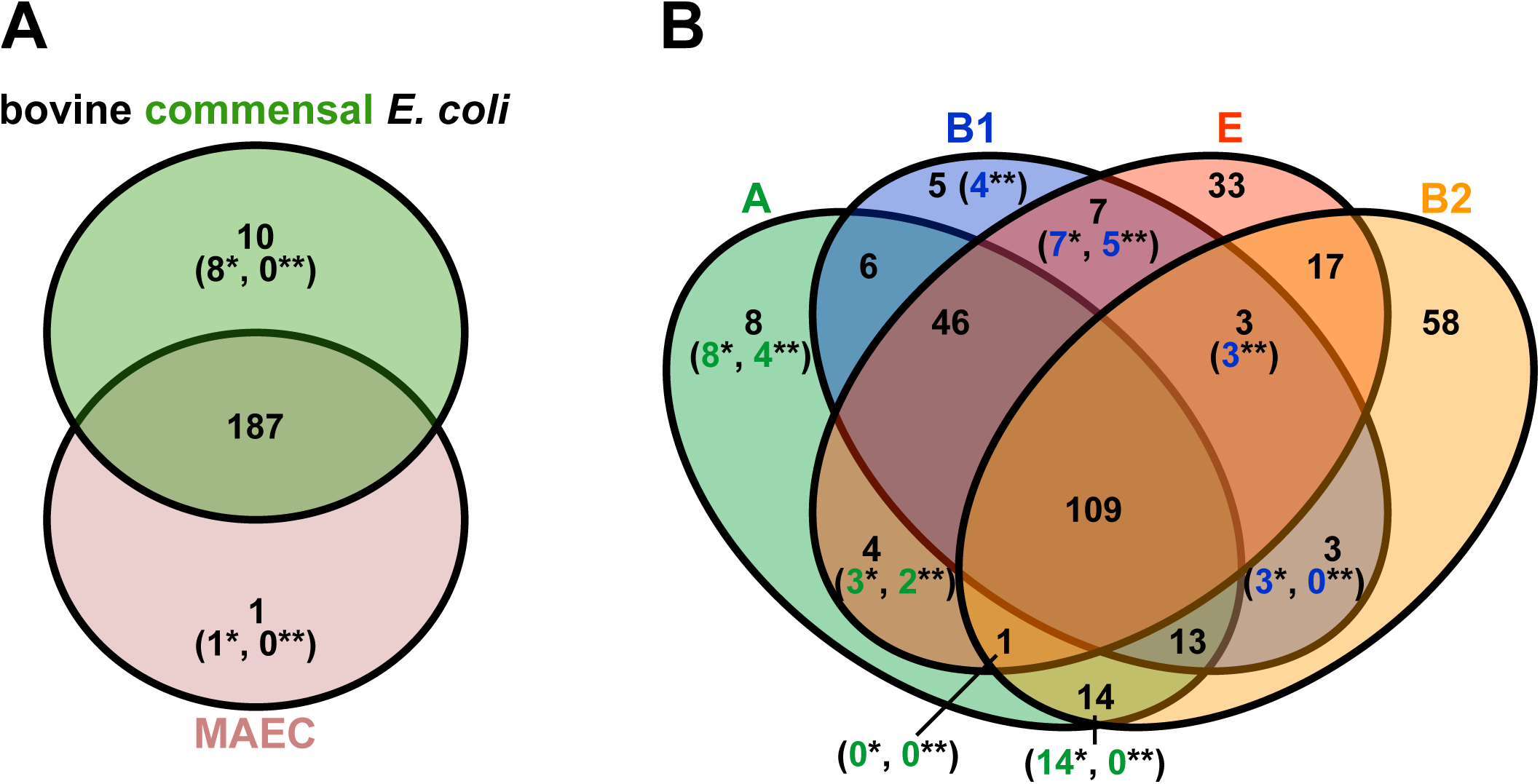
Venn diagrams of virulence-associated gene enrichment in pathotypes and phylogroups. Enriched virulence-associated genes (numbers without parentheses) were identified with 70% inclusion and 30% exclusion group cutoffs for the bovine-associated *E. coli* classified either by A) pathotype (MAEC or commensal) or B) phylogenetic groups (A, B1, B2, or E). Statistical significance of VF association was tested with Fisher’s exact test (p < 0.05, numbers with a single asterisk) and Bonferroni corrected (numbers with two asterisks). In the phylogroups association was only tested for the multi-genome phylogroups A versus B1. As with the OG enrichment analysis, phylogenetic lineage of the strains dominates VF content and only very few virulence-associated genes were enriched in the pathotypes.

Because T3SS-related genes were present in MAEC and commensals, we wanted to analyze the ETT2 determinant in more detail in our strain panel. In addition to ETT2, we also examined the large ECC-1470 T6SS/1 and Flag-2 gene regions. All three putative virulence regions show a high amount of mutational isoforms and/or absence in the strain panel (Figure 4), warranting a detailed analysis. For this purpose, the gene composition of such regions was depicted for all bovine-associated *E. coli* from the strain panel (Figure 6 and Additional file 3: Figure S7A and B). In the case of strain D6-117.29 the ETT2 and T6SS regions could probably not be fully manually assembled, because of the high fragmentation of the genome.

**Table 2.** **Virulence-/fitness-associated genes significantly associated in MAEC or commensal isolates, as well as phylogroups A or B1**

| Gene/locus tag | Accession number | VF class | Phylogroup association, enrichment |
| --- | --- | --- | --- |
| **MAEC-enriched virulence-/fitness-associated gene** |  |  |  |
| *eprI* | YP_006097353 | T3SS/ETT2 | significantly A-associated |
| **Commensal-enriched virulence-/fitness-associated genes** |  |  |  |
| EC042_1639 | YP_006095949 | CU fimbriae | significantly B1-associated |
| *ydeT* | YP_006095947 | CU fimbriae | significantly B1-associated |
| *iucA* | NP_755502 | Iron uptake | no hit |
| *iucB* | NP_755501 | Iron uptake | no hit |
| *iucC* | NP_755500 | Iron uptake | no hit |
| *iucD* | NP_755499 | Iron uptake | no hit |
| *iutA* | NP_755498 | Iron uptake | no hit |
| *shiF* | NP_755503 | Iron uptake | no hit |

MAEC, mastitis-associated *E. coli*; CU, chaperone usher; T3SS, type III secretion system; ETT2, *E. coli* T3SS 2

The ETT2 gene cluster shows high genetic flexibility and many deletions and insertions (Figure 6). Nevertheless, small features still reveal a phylogenetic relationship of similar pseudogene composition. For example *eprJI*, *orgB*, and *epaO* are mostly pseudogenes in B2 strains, but the genes seem to be functional in all phylogroup A and E strains. No comparable pattern was found in relation to the pathotypes. Almost all genomes lack a fragment present in the putatively intact ETT2 region of phylogroup D1 EAEC strain 042 (*eivJICAEGF*), which is located between two small direct repeats and thus often deleted [43]. Only the ETT2 gene cluster in phylogroup E isolate D6-113.11 has an identical structure as 042 (phylogroup E is most closely related to phylogroup D1).

The Flag-2 region is basically present or entirely absent in the strain panel. No intermediate attrition isoforms are observable (Additional file 3: Figure S7A). A large deletion is apparent in O157:H43 strain T22. This deletion encompasses the whole Flag-2 region and respective flanking backbone genes. Thus, *E. coli* O157:H43 strain T22 was omitted from the diagram. Additionally, the deletion includes also the housekeeping genes downstream of the T6SS/1 gene cluster of *E. coli* ECC-1470 indicated by dots in Additional file 3: Figure S7B.

The subtype i1 T6SS/1 of MAEC strain ECC-1470 is the most variable of the virulence-associated regions investigated in more detail in this study, with many repetitive sequence subregions. Typical for T6SSs, it is also adjacent to a highly repetitive *rhs* element. Strain ECA-727 lacks the *yafT* to *impA* genes, because of a putative phage insertion in this region. This phage is not included in the figure and the truncation is indicated by dots in the diagram. The T6SS determinants in MAEC strains 1303, MPEC4839, D6-117.29, D6-113.11, and commensal RiKo 2305/09 are most likely not functional because of their small sizes. Overall, we could not find any features of this gene cluster which are associated with phylogeny or pathogenicity of the strains.

**Figure 6.**
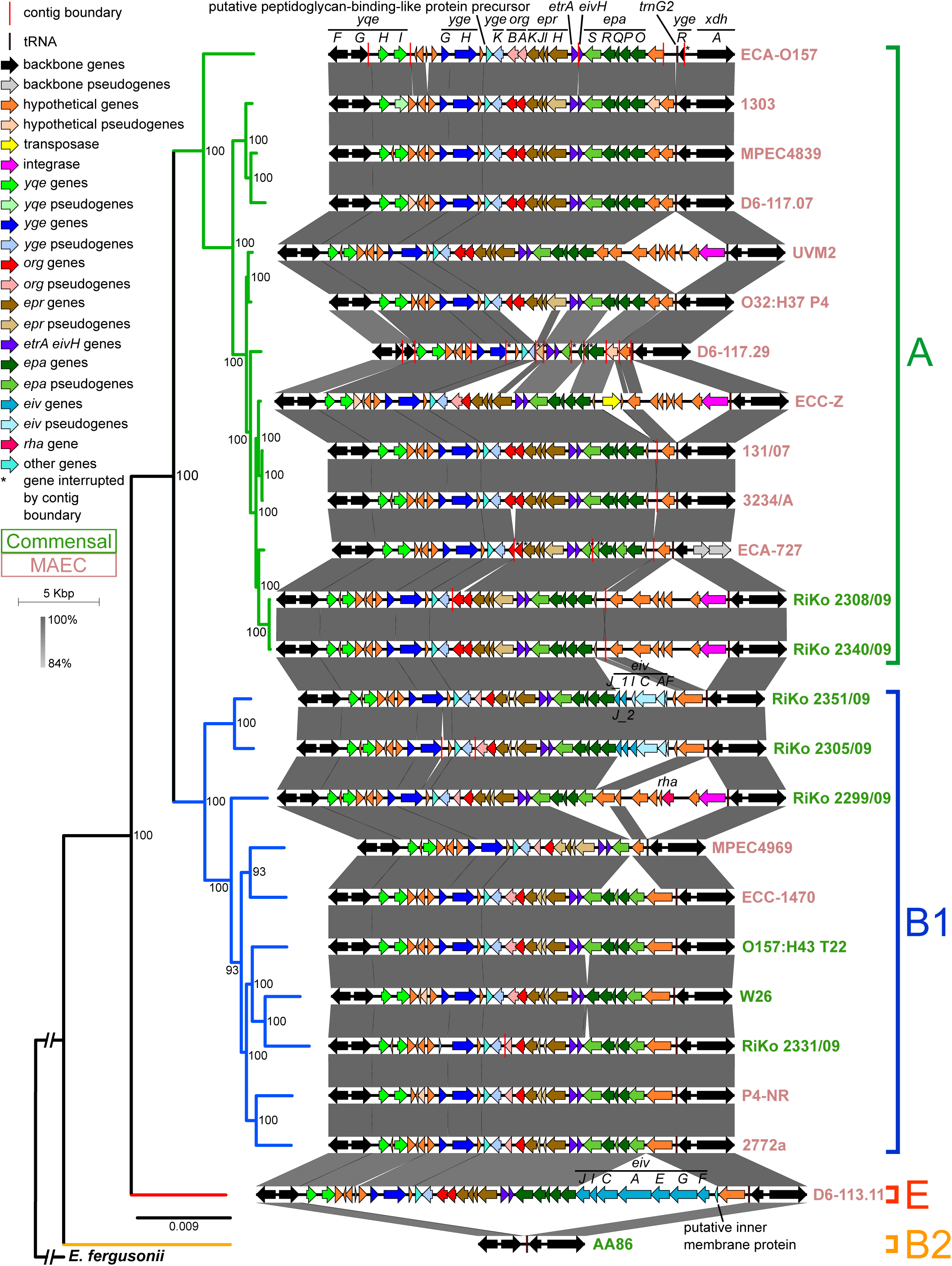
Gene organization of the ETT2 gene cluster in the bovine-associated *E. coli* genomes. Comparison of the ETT2 gene cluster in the *E. coli* of the strain panel based on BLASTN+. Homologous regions are connected via grey vertices and colored by nucleotide identity. The genomes are ordered according to the WGA core genome phylogeny (Additional file 3: Figure S2A), which is attached on the left side (bootstrap support values below 50 were removed). Phylogroups are indicated correspondingly. MAEC strain names are colored in light red and commensals in green. Gene names are indicated above genomes encoding for these. The respective contigs of the draft genomes containing the gene cluster were concatenated (contig boundaries are indicated by red vertical lines) and CDS spanning contig borders reannotated if needed (indicated by asterisks). ETT2 contigs of genome D6-117.29 were difficult to concatenate, because of its high fragmentation. Backbone genes not belonging to ETT2 are colored black. Genes within the ETT2 region have different colors (see the legend) to be able to evaluate their presence. Pseudogenes have a lighter color fill. ETT2 shows a large number of different mutational isoforms. Nevertheless, ETT2 composition follows phylogenetic history rather than pathotype affiliation.

## Discussion

This is the first study which investigates *E. coli* genomes in relation to bovine mastitis including two closed genomes of finished quality [44], MAEC 1303 and ECC-1470 [45]. Closed genomes of a finished quality allow insights into the genome organization, synteny, and detection of MGEs. We additionally sequenced six bovine fecal commensal and six MAEC draft genomes and supplemented these with publicly available reference bovine *E. coli* strains. With this strain panel of 16 bovine mastitis and nine bovine *E. coli* commensal isolates we were able to analyze differences in the gene content between MAEC and commensal strains in relation to the phylogenetic as well as genomic diversity of bovine *E. coli* in general. Bovine strains are phylogenetically diverse and do not show a virulence-related gene content that is associated with either pathotype. This has implications for the definition of mastitis-related VFs and a bovine mastitis *E. coli* pathotype.

The assembly statistics of the draft genomes of this study indicate a suitable quality for the purposes of our analyses, with 24 to 290 contigs and N50 values ranging from 79 to 358 kb for contigs larger than 500 bp (Additional file 5: Table S3). There are four apparent exceptions: First, the genome of commensal reference strain AA86 has gone through multiple gap closure steps and has only five contigs with an N50 of 2,860 kb [46]. Two of these five contigs are plasmids, making AA86 the only strain with resolved plasmid sequences in the strain panel in conjunction with the finished 1303 and ECC-1470 genomes [45]. Second, the three MAEC reference draft genomes D6-117.29, ECA-727, and ECA-O157 are highly fragmented with more than 500 contigs each. However, their coding percentage and overall CDS numbers are in the range of other *E. coli* genomes and thus they were included in the strain panel (Additional file 1: Table S1). Also, overall presence of VFs in the strain panel did not relate to contig number (Additional file 14: Dataset S7).

### Bovine-associated *E. coli* originate mostly from phylogroups A and B1

*E. coli* phylogroup A is traditionally associated with commensal strains, while its sister taxon B1 is associated with commensals and different IPEC including ETEC, EAEC, and EHEC [42, 17, 18]. ECOR phylogroup E includes the genetically closely related O157:H7 EHEC and O55:H7 EPEC [47]. Interestingly, the bovine commensal O157:H43 isolate T22, even though belonging to the O157 serotype, is not a member of phylogroup E, but of group B1 [48], providing an example for horizontal transfer of O-antigen genes. Finally, phylogroup B2 is the most diverse phylogroup, based on nucleotide and gene content. This group also includes most of the ExPEC, like UPEC, APEC, and MNEC [42, 49, 17, 12]. However, with the accumulation of *E. coli* sequencing data, the traditional association of phylogroups with pathotypes have softened, as many pathotypes were shown to have emerged in parallel in different lineages [14, 42, 18]. The phylogenetic placement of the bovine isolates used in this study is in agreement with previous studies where MAEC and bovine commensals were also enriched in phylogroups A and B1, while other phylogroups play only a minor role [11, 9]. Depending on the study and the respective analyzed strain panel, MAEC isolates are either more common in phylogroup A [6, 22, 23] or phylogroup B1 [11, 27, 50]. Also, the WGA phylogeny shows that bovine MAEC and commensals do not cluster together, but rather originate from diverse lineages within phylogroups (Figure 1) [11, 26, 27]. The discrepancies of MAEC phylogroup associations between the previous studies might be a result of country-specific differences or differences in sampling and phylotyping techniques. The polyphyletic evolutionary history of bovine *E. coli* (both MAEC or commensals) is substantiated by their high genotypic and phenotypic plasticity [11, 6, 9, 24, 5]. In light of these studies and the genealogy of the bovine-associated *E. coli* in this work (Figure 1) the strain panel is suitable and sufficiently diverse in its phylogeny for more detailed comparative analyses of MAEC and commensal bovine *E. coli* genomes. Two possible explanations for the phylogenetic diversity of MAEC and bovine commensals can be considered. On the one hand, the ability to cause mastitis could have been developed in parallel on several independent occasions during the evolutionary history of *E. coli* by selecting forces [26]. On the other hand, MAEC might be recruited from the normal intestinal commensal microbiota and the ability to cause mastitis is facultative, as has been proposed for ExPEC [19, 18, 12, 51].

### Gene content of bovine-associated *E. coli* mirrors phylogeny rather than pathotype

Several studies have shown that recombination between extant *E. coli* phylogroups is limited by phylogenetic diversity [29, 30]. Thus, the phylogenetic background of *E. coli* has a big impact on possible recombination events and most importantly on the gene content of the flexible genome [18, 15]. Nevertheless, there are examples of convergent evolution in *E. coli,* especially in IPEC pathotypes from multiple parallel phylogenetic origins that typically contain a specific set of VFs, e.g. the occurrence of EHEC in the distant phylogroups B1 and E mediated by HGT of MGEs [47, 29]. Our study demonstrates that there is no evidence for HGT of large mastitis-specific genomic regions, and that the phylogenetic background of the strains has a deciding impact on the overall gene content (Figure 2 and Additional File 3: Figure S2B). A clustering of strains according to pathotypes would have hinted towards a common gene content and a difference in ecological lifestyles and habitats, as a result of positive selection on the ancestral genomes [29]. However, our results demonstrate that the flexible and the core genome appear to coevolve.

Two previous studies with similar methodology came to two different conclusions. Blum *et al.* [25] reasoned that three mastitis strains (O32:H37 P4, VL2874, VL2732) were much more closely related in gene content compared to an environmental (commensal fecal) strain (K71), based on the different pathotypes. However, the MAEC in this study are phylogenetically strongly related (phylogroup A) whereas the single commensal strain belongs to phylogroup B1. Thus, as we observed in our study, the phylogenetic relationship had a strong impact on the gene content dendrogram. Kempf *et al.* [27] comparing four phylogroup A MAEC (D6-117.07, O32:H37 P4, VL2874, VL2732), one phylogroup E MAEC (D6-113.11), and the K71 commensal, achieved results comparable to ours. The authors argued that mastitis pathogens with different phylogenetic histories might employ different virulence strategies to cause mastitis, similar to the variable VF repertoire of ExPEC. A hypothesis we tested in this study by searching for well-known *E. coli* VFs in the bovine-associated *E. coli* strains discussed below.

### MAEC cannot be distinguished from bovine commensal *E. coli* based on the presence of virulence-associated genes

It was suggested that the genome content of MAEC is distinct from bovine commensals and not random, as a result of selective pressure. VFs important for MAEC pathogenicity would then supposedly be positively selected within the bovine udder [25, 26]. Several virulence-associated properties have been proposed for MAEC pathogenicity [52, 27, 3]: multiplication and persistence in milk and the udder [10, 53], resistance to serum components and neutrophil neutralization mechanisms [54, 7, 55], adhesion to (and invasion of) mammary epithelial cells [10, 4, 6, 56], and stimulation of the innate immune response by PAMPs [57, 58]. Against this background, a myriad of previous publications have tried to identify VFs specific for MAEC with varying degrees of success [11, 6, 59, 22, 60, 55, 61, 50, 23, 24]. However, the results of these studies do not agree upon the identified VFs, which is due to the diversity of MAEC and bovine *E. coli*, generally. The aforementioned publications followed a classical diagnostic typing procedure by using PCR assays for virulence-associated gene detection. Only Kempf and colleagues applied a bioinformatic approach similar to ours with a candidate VF panel of 302 genes [27]. However, our larger strain and selected VF panels enabled a more detailed analysis. In our study, commensal strains and MAEC exhibited a similar average virulence-associated gene presence (250 and 237, respectively), also comparable to averages of the *E. coli* genomes categorized by the different phylogroups (phylogroup A: 233, B1: 254, B2: 227, and E: 235). Additionally, we used a bottom-up approach to identify overall OGs associated with MAEC. For these analyses we allocated the analyzed strains into pathotypes (MAEC or commensal) based on their source of isolation and used 70%/30% inclusion/exclusion cutoffs to detect OG/VF association with either group. Furthermore, we applied Fisher’s exact test to determine statistically significant associations between OGs or VFs and pathotypes or phylogroups. Both our comparisons of the association of OGs and VFs with mastitis or commensal genomes did not reveal a significant correlation between the presence of individual virulence-related genes and the mastitis-associated isolates (Figures 3A and 5A, Additional file 7: Dataset S3 and Additional file 16: Dataset S8). Using 70%/30% inclusion/exclusion cutoffs, we recovered only one significantly MAEC-and eight significantly commensal-enriched VF genes (Table 2). As expected from the OG analysis (Figure 3B and Additional file 8: Dataset S4), the phylogroups had also a strong impact on VF enrichment (Figure 5B and Additional file 17: Dataset S9).

The aerobactin gene cluster together with the *iutA* and *shiF* genes were detected as significantly associated and enriched in the commensal strains in both our OG and VF analyses (Table 2 and Additional file 8: Dataset S4). The siderophore system aerobactin is considered an ExPEC VF needed for iron uptake under limiting conditions, e.g. in the urinary tract or serum [62]. These genes are often encoded by plasmids harboring additional traits, like colicins and other iron transport systems, e.g. in APEC colicin plasmids [62, 63]. Thus, its distribution might also be due to positive selection of beneficial traits for commensalism, which are encoded by the same plasmid. Another group of commensal-enriched virulence genes included fimbriae-associated genes (EC042_1639, *ydeT*) (Table 2). However, these two genes have in common, that they are enriched in phylogroup B1.

The *eprI* gene was determined as the only significantly mastitis-associated and -enriched virulence-associated gene (Table 2). This gene, which was also shown to be significantly associated with phylogroup A strains, belongs to the ETT2 determinant, a large gene cluster with frequent deletion isoforms in *E. coli [43]*. The ETT2 type III secretion system is contained on GI8 of *E. coli* 1303 and on GI9 of strain ECC-1470 (Additional file 10: Dataset S5 and Additional file 11: Dataset S6). ETT2 has not only been discussed as a VF during mastitis [64], but has also been implicated in being involved in invasion and intracellular survival of blood-brain barrier cells of MNEC K1 strains [65] and in serum resistance of APEC O78:H19 strain 789, besides its degenerate form in the strain [62]. Its prevalence has been analyzed in bovine mastitis *E. coli* isolates and was determined to be approximately 50% [64]. ETT2 has different mutational attrition isoforms in our bovine-associated strain panel, supporting the results of an earlier study [64]. However, overall ETT2 presence was not related to MAEC (Figure 6). Based on the comparative analysis, and in accordance with Blum *et al.* [10] these results suggest that serum resistance is not an essential trait for the ability of MAEC to cause intramammary infections. Thus, a role of ETT2 in MAEC is debatable, especially since only *eprI* and none of the other ETT2 genes were MAEC-enriched. In conclusion, MAEC are characterized by a lack of “bona fide” VFs [11, 27, 24]. Instead, the VF variety observed rather mirrors the genome plasticity of bovine-associated *E. coli*, regardless of pathotype. Although many of these putative VFs are not connected to mastitis virulence, they are still maintained within the genomes. This suggests that they serve as FFs for gastrointestinal colonization and propagation, rather than VFs.

### Large virulence regions and intraphylogroup comparisons of putative VFs are also not pathotype-specific

An alternative flagellar system (Flag-2) is encoded on 1303 GI1 [43]. The Flag-2 locus encodes also for a type III secretion system in addition to the alternative flagellar system, which might be in cross-talk with ETT2. In contrast to the typical *E. coli* peritrichous flagella 1 gene cluster (Flag-1), which is a polar system for swimming in the liquid phase, the lateral Flag-2 most likely has its functionality in swarming ability over solid surfaces [43]. Flagella are important for motility, but also for adherence during host colonization and biofilm formation [16]. Additionally, flagella might play an important role in the udder for dissemination from the teat and counteracting washing out during milking [10]. MAEC ECC-1470 also carries two T6SS determinants located on GI1 (designated as the first ECC-1470 T6SS, T6SS/1) and on GI8 (ECC-1470 T6SS/2), respectively. *E. coli* ECC-1470 T6SS/1 is classified as subtype i1 [66] or the second *E. coli* T6SS-2 phylogenetic cluster [67] and T6SS/2 as subtype i2 [66] or the first *E. coli* T6SS-1 cluster [67]. Subtype i1 T6SSs generally participate in interbacterial competition, subtype i2 T6SSs target eukaryotic cells and play a role in the infection process of pathogens. All T6SS are implicated in mediating adherence and biofilm formation [67]. The GI1-encoded T6SS/1 was consistently present in strains ECA-O157, ECA-727, and ECC-Z, but only sporadically in human reference commensal strains, and thus associated with MAEC in a preceding study [28]. Nevertheless, the corresponding phenotypes of these systems are mainly unknown and their function, especially any putative role in mastitis, might well be indirect [67]. Because of a low prevalence of T6SS genes in the five included MAEC genomes and presence in commensal strain K71, another previous study questioned the role of T6SS systems in mastitis [27]. We can support this study, as even our detailed analysis of the ETT2, Flag-2, and T6SS/1 regions did not reveal any association with MAEC isolates in our strain panel. These regions of the flexible genome mirror the underlying genomic and phylogenetic diversity of bovine *E. coli*.

Several studies argue that MAEC strains from divergent phylogenetic backgrounds might use different VF subsets and virulence strategies to elicit bovine mastitis [10, 22, 26, 27]. We tested this hypothesis exemplarily, by analyzing the 31 genes of the *fec*, *paa*, and *pga* regions for pathotype enrichment within the multi-genome phylogroups of our strain panel, A and B1. These three regions were detected as being essential in phylogroup A MAEC [26], but they have not been analyzed in other phylogroups. Five of the *pga* and all *fec* genes were significantly associated with MAEC, however *fecBCDE* also with phylogroup A (Additional file 18: Dataset S10). Also, all genes of the three regions were included in the all-strain soft core with the 70% inclusion threshold (Additional file 15: Table S7). Thus, none of the genes were associated with pathotype and only the *paa* phenylacetic acid degradation pathway determinant was missing in the single-genome ECOR phylogroups B2 and E. This might have tipped the scales in the analysis of phylogroup A genomes by Goldstone and co-workers. The 13 phylogroup A strains of our strain panel contain eleven MAEC and two commensal isolates. The ten strains of phylogenetic lineage B1 comprise four MAEC and six commensal strains. Due to ongoing sequencing efforts, the number of suitable reference genomes for more detailed analyses is likely to increase in the near future. However, this is the first study to be able to perform such an analysis. In the ECOR group A genomes, the *fec*, *paa*, and *pga* regions were not pathotype-enriched (with the 70%/30% inclusion/exclusion cutoffs), but were present in the group soft core (except for *paaB* in the unspecific category; Additional file 19: Dataset S11) and none were statistically significantly associated via Fisher’s exact test. In a similar way, the PGA biosynthesis and Fec-system encoding regions were also mostly categorized into the group soft core of the analysis with B1 strains (Additional file 20: Dataset S12) and again with no significant Fisher’s exact test p-values. Only *fecBCDE* were in the unspecific category, because these genes are missing in the genomes of the commensal isolates RiKo 2331/09, O157:H43 T22, and W26. However, the whole seven-gene *pga* region was MAEC-enriched in our phylogroup B1 strain set (albeit without significance), present in all four MAEC, but only in two of the six commensals. We want to stress that this result depends highly on the strain collection used and more bovine *E. coli* strains, especially commensals, from all available phylogroups need to be incorporated for an in-depth analysis. As all three regions are present in the all-strain soft core genome of our whole strain panel analysis, these results illustrate the drawbacks of inferring general observations from low numbers of strains (especially when focusing only on pathogenic strains) considering the genome plasticity of bovine *E. coli*.

## Conclusions

This is the first publication to include a phylogenetically diverse bovine *E. coli* strain panel incorporating both MAEC and commensal isolates for genomic content comparisons. Besides the two closed bovine MAEC 1303 and ECC-1470 [45], that can serve as high sequence and annotation quality references, this study includes the largest collection of bovine *E. coli* commensals from fecal origin of udder-healthy cows [68]. As we could not identify any genes significantly associated with MAEC that were not also present in commensal strains or correlated with the strains’ phylogenetic background, an MPEC pathotype characterized by specific VFs could not be defined. It is more likely that virulence-associated genes, which have been previously implicated in facilitating mastitis, have their principal function in colonization and persistence of the gastrointestinal habitat. Thus, like ExPEC, MAEC are facultative and opportunistic pathogens basically of naturally occurring commensal (“environmental”) *E. coli* origin [19, 18, 23, 12, 51, 24]. As a consequence, we propose to use the term mastitis-associated *E. coli* (MAEC) instead of mammary pathogenic *E. coli* (MPEC).

The genome content of certain bovine *E. coli* strains seems not to support the ability to elicit mastitis in udder-healthy cows as was shown in the case of the commensal strain K71 [25]. The large pan-genome of bovine *E. coli* isolates offers many gene combinations to increase bacterial fitness by utilization of milk nutrients and evasion from the bovine innate immune system, thus resulting in sufficient bacterial intra-mammary growth and consequently infection of the mammary gland [10, 53, 69, 70]. Isolates with an increased potential to cause mastitis can colonize the udder by chance depending on suitable environmental conditions and the cow’s immune status. Our data also demonstrate, that there is no positive selection in MAEC for the presence of virulence-associated genes required for causing mastitis. This has implications for vaccine development and diagnostics. Reverse vaccinology may not be suitable for the identification of specific MAEC vaccine candidates, and the utilization of marker genes for improved diagnostics and prediction of the severity and outcome of an *E. coli* bovine mastitis might fail. Herd management and hygiene are still the two most important factors for preventing *E. coli* mastitis incidents. Several studies have shown a dramatic decrease in the bovine udder microbiome during mastitis, even after recovery [71-73]. It might be worthwhile to consider alternative prevention strategies like strengthening the natural udder microbiota that competes with pathogens [74].

We urge the research community to not fall into the same trap with whole genome studies as with the previous typing studies. Mastitis researchers need to consolidate their efforts and, as Zadoks *et al.* eloquently put it, not to waste precious resources on “YATS” (yet another typing study) [5]. It is necessary to step away from the reductionist approach and adapt an integrated course of action by examining the host-pathogen interaction simultaneously. Synergistic application of techniques, like dual RNA-Seq of host and bacteria [75], Tn-Seq to test virulence association of genes *in vivo*, comparative SNP analysis of orthologous genes and intergenic regions, proteomics, and metabolomics, are readily available to correlate physiological traits with genomic information. Additionally, the comparison of closed genomes offers the possibility to comprehensively analyze the complete genomic context of strains including genomic architecture, rearrangements and movement of mobile genetic elements.

## Methods

A detailed methods section can be found in Additional File 21.

### Bacterial strains, isolation, and published reference genome acquisition

All fourteen isolates in this study were collected using routine clinical practices from the bovine habitat. Commensal strains were isolated from fecal samples of udder-healthy and mastitis-associated strains from the serous udder exudate of mastitis-afflicted cows. Mastitis strains were acquired from different veterinary diagnostic laboratories in the indicated countries, listed in the genomes feature overview table (Additional file 1: Table S1). Additionally, eleven draft bovine-associated *E. coli* reference genomes were downloaded from NCBI to be used in the analyses. See Table 1 for the respective reference publications. The corresponding accession numbers are given in Additional file 1: Table S1.

### Library preparation and sequencing

The strains with closed genomes, 1303 and ECC-1470, were sequenced on a 454 Titanium FLX genome sequencer with GS20 chemistry as described in [45].

These two strains were additionally and the draft strains [68] solely sequenced with a 101-bp PE sequencing run on a HiScan SQ sequencer (Illumina, San Diego, CA, USA).

### Assembly of the genomes

Both 454 read sets for the genomes of *E. coli* 1303 and ECC_1470 were *de novo* assembled with Newbler (Roche) (v2.0.00.20 for 1303 and v2.3 for ECC-1470) [76]. Additionally, these reads were assembled in a hybrid *de novo* approach in combination with the respective Illumina reads using MIRA (v3.4.0.1) [77]. Afterwards, each 454 Newbler assembly was combined with the respective hybrid assembly in Gap4 (v4.11.2) of the Staden software package [78]. The remaining gaps in the assembly were closed by primer walking via directed PCR and Sanger sequencing utilizing BigDye Terminator chemistry with ABI 3730 capillary sequencers. The closed genomes were edited to the “finished” standard [44].

The Illumina reads from the draft *E. coli* genomes were each randomly subsampled to an approximate 70-fold coverage with seqtk (v1.0-r32; https://github.com/lh3/seqtk). Afterwards, the PE reads were *de novo* assembled with SPAdes (v3.1.1) [79] and only contigs >= 500 bp retained. At last, the assembled contigs were ordered against the respective *E. coli* 1303 or *E. coli* ECC-1470 reference genomes, according to the ECOR phylogroup affiliation of the draft genomes. All Sequence Read Archive (SRA) study accession numbers for the Illumina and 454 raw reads of the *E. coli* genomes of this study can be found in Additional file 5: Table S3. This file also includes the assembly statistics for all 23 bovine-associated *E. coli* draft genomes. The draft genomes of this study are in the “high-quality draft” standard [44].

### Annotation of the genomes

The complete genome sequences of *E. coli* 1303 and ECC-1470 were initially automatically annotated with Prokka (v1.9) [80] and the annotations subsequently supplemented with further databases. This automatic annotation was manually curated with Artemis (v15.1.1) [81] and tbl2tab (v0.1) [82]. Additionally, the annotations of *E. coli* strains 1303 and ECC-1470 were compared (ACT, v12.1.1 [83], and BLASTN+, v2.2.28 [84]) and adapted to each other for a uniform annotation. The high quality annotation of the *E. coli* 1303 genome was then used as reference for the ECOR phylogroup A strains and the ECC-1470 genome annotation for the ECOR B1 strains during the Prokka annotation of the 12 draft genomes of this study.

All eleven reference strains were also automatically reannotated with Prokka to have a uniform ORF-finding and facilitate comparative genomics. The annotations of the references were shortly manually curated in the three putative virulence regions ETT2, Flag-2, and strain ECC-1470’s T6SS/1 by comparisons to the 1303 and ECC-1470 genomes as mentioned above. GENBANK files for these reannotations can be found in Additional file 22: Dataset S13 and Additional file 23: Dataset S14. For an overview of the annotations see the genome feature table created with genomes_feature_table (v0.5) [82] (Additional file 1: Table S1). This table also includes the reference *E. coli* genomes for the phylogenetic analysis (see below), however their annotation features are listed as downloaded from NCBI.

### Phylogenetic analysis

A WGA of selected *E. coli* genomes was done with the default parameter settings of Mugsy (v1.2.3) [85] and with *E. fergusonii* as outgroup. The MAF alignment file was further processed to contain only locally colinear blocks without gaps present in all aligned genomes utilizing the software suite Phylomark (v1.3) [86]. The concatenated and filtered alignment was then subjected to RAxML (v8.1.22) [87] to infer the best scoring ML phylogeny with 1,000 bootstrap resamplings for local support values. The resulting tree was visualized with Dendroscope (v3.4.4) [88]. This phylogeny was used to classify the bovine-associated strains into ECOR phylogroups according to the included reference strains (with a known phylogeny) and monophyletic clades. The same procedure was followed including only the 25 bovine-associated *E. coli* strains. This tree was visualized with FigTree (v1.4.1; http://tree.bio.ed.ac.uk/software/figtree/) midpoint rooted.

STs were assigned with ecoli_mlst (v0.3) [82] according to the Achtman *E. coli* MLST scheme [13] employing NUCmer with default parameters. PHYLOViZ (v1.1) [89] was used to create a MST with the goeBURST algorithm [90] to classify the STs into CCs. CC numbers were allocated according to the Achtman *E. coli* MLST database. All allele, ST, and CC numbers can be found in Additional file 2: Table S2.

### Detection of genomic islands and prophages, and generation of circular genome diagrams

GIs and prophages were predicted in the two closed genomes. GIs were predicted with the three prediction methods of IslandViewer 3 [91]: the two sequence composition methods SIGI-HMM [92] and IslandPath-DIMOB, and the comparative genomic prediction method IslandPick [93]. Only predicted GIs with a size greater than 8 kb were retained. Prophages were predicted with the PHAge Search Tool (PHAST) [94]. Circular genome views were created with the BLAST Ring Image Generator (BRIG, v0.95) using BLASTP+ (v2.2.28) [84] with a disabled low complexity filter (option ‘-seg no’) and upper/lower identity thresholds set to 90% and 70%, respectively. The location of the predicted GIs and prophages are visualized in these diagrams.

### Identifying serotypes

The SerotypeFinder (v1.0) database was used to determine serotypes *in silico [95]*. For some strains SerotypeFinder could not resolve the O-or H-antigen uniquely, in these cases both are listed.

### Ortholog/paralog analysis

Orthologous and paralogous proteins in all 25 bovine-associated genomes were identified with Proteinortho (v5.11) [96, 97] with a 1 x 10^−5^ E-value and 70% coverage/identity cutoffs. This resulted in a total number of 13,481 OGs from the overall 116,535 CDSs in the bovine-associated strain panel.

To identify pathotype-(mastitis/commensal) or phylogroup-enriched (ECOR phylogroups A/B1/B2/E) OGs we employed Fisher’s Exact test provided in R (v3.2.5) and tested for OGs which are significantly (p<0.05 and with a Bonferroni correction) associated with the different pathotype or phylogenetic groups. Additionally, OGs were considered enriched if they are minimally present in 70% of the genomes of one genome group (inclusion cutoff) and in maximally 30% of genomes of all other groups (exclusion cutoff) using po2group_stats (v0.1.1) [82]. The Fisher’s exact test p-values were visualized as Manhattan plots with R package ggplot2 (v2.2.0) [98] (Additional File 4: Dataset S1). An “all-strain soft core genome” over all genomes with the 70% inclusion cutoff (rounded 18 genomes of the total 25) was determined.

The resulting pathotype-enriched OGs were further evaluated by comparing their representative proteins to the representative proteins in the phylogroup-enriched categories and the all-strain soft core. For this purpose, the prot_finder pipeline with BLASTP+ was used, as described below in the VF workflow, with the pathotype-enriched representative proteins as queries and the phylogroup-enriched or all-strain/phylogroup soft core proteins as subjects.

Finally, a gene content tree was calculated using RAxML (v8.0.26) and 1,000 resamplings with the Proteinortho presence/absence matrix of OGs (included in Additional File 4: Dataset S1).The clustering tree was visualized midpoint rooted with Figtree.

### Screening of the genomes for known virulence factors

VF reference protein sequences were collected from the VFDB ([99-101] and the primary literature. For an overview of the VF panel see Additional file 12: Table S5 and the GitHub repository https://github.com/aleimba/ecoli_VF_collection (v0.1) [41].

The VF panel was used to assess the presence/absence of the 1,069 virulence-associated genes in the annotated bovine-associated strains with the prot_finder pipeline (v0.7.1) [82] using BLASTP+ (v2.2.29) with the following options: 1 x 10^−10^ E-value cutoff (‘-evalue 1e-10’), 70% query identity and coverage cutoffs (options ‘-i’ and ‘-cov_q’), and the best BLASTP hits option (‘-b’). As with the gene content tree, a ML RAxML BINGAMMA search was done to cluster the results in the VF binary matrix with 1,000 resamplings. Additionally, the binary VF hit matrix was visualized with function heatmap.2 of the R package gplots and R package RColorBrewer (v1.1-2) [102]. The RAxML cladogram was attached to this heatmap with R package ape (v3.4) [103]. The binary matrix, the cladogram NEWICK file, and the R script are included in Additional file 14: Dataset S7.

VF associations with either pathotypes or phylogenetic groups were tested with a two-tailed Fisher’s exact test for significance (p < 0.05) and with a Bonferroni-corrected significance value. Manhattan plots were created with R package ggplot2 (Additional File 4: Dataset S1). Again, inclusion and exclusion cutoffs were set to 70% and 30%, respectively, using binary_group_stats (v0.1) [82]. Also, an all-strain soft core VF set was calculated over the virulence-associated gene hits of all genomes with a 70% (18 genome) inclusion cutoff. Pathotype-enriched VF proteins were compared to phylogroup-enriched VF proteins for evaluation.

The same prot_finder pipeline and binary_groups_stats workflow was also used for two putative MAEC-specific regions in ECOR phylogroup A genomes [26], which are not included in the VF panel. The first region is the biofilm-associated polysaccharide synthesis locus (*pgaABCD-ycdT-ymdE-ycdU*). The locus tags in *E. coli* genome 1303 are EC1303_c10400 to EC1303_c10440, EC1303_c10470, and EC1303_c10480. The second region encodes proteins involved in the phenylacetic acid degradation pathway (*feaRB-tynA-paaZABCDEFGHIJKXY*; MG1655 locus tags b1384 to b1400). The third region (the Fec uptake system, *fecIRABCDE*) is already included in the VF panel of this study. For this analysis the resulting binary BLASTP+ hit matrix was also tested with binary_groups_stats for pathotype association within the ECOR A and B1 phylogroups of the bovine-associated strain panel (with the 70% inclusion and 30% exclusion cutoffs). Associations were additionally tested with Fisher’s exact test for significance.

### Analysis of large structural putative virulence regions

The composition of the large virulence regions ETT2, Flag-2, and the T6SS/1 subtype i1 determinant of *E. coli* ECC-1470 as well as the antimicrobial multidrug resistance element of 1303 (AMR-SSuT in GI4) was compared in more detail for the bovine-associated strain panel with Easyfig (v2.2.2) [104].

### General data generation and figure editing

Dendroscope was used to create tanglegrams between the cladograms of the bovine-associated strain panel WGA phylogeny, gene content, or VF clustering trees. All figures were created either with R (v3.2.5) [105] for the heatmap, Manhattan plots, or venn diagrams, Dendroscope or FigTree for phylogenetic trees, PHYLOViZ for the MLST MST, or Easyfig for the genome diagrams and color edited, labelled, or scaled with Inkscape (v0.91) without changing data representation. The only exception are the BRIG circular genome diagrams which were edited with Gimp (v2.8.16).

## Additional files

**Additional file 1: Table S1.** Genome feature table for the 64 *E. coli*, four *Shigella* spp., and the one *Escherichia fergusonii* genomes plus accession numbers. (XSLX 18 KB)

**Additional file 2: Table S2.** MLST allele profiles, ST and CC numbers for the 64 *E. coli* and four *Shigella* spp. strains. (XSLX 10 KB)

**Additional file 3: Figure S1.** Minimum spanning tree (MST) of the MLST results. **Figure S2.** Phylograms and tanglegrams for the 25 bovine-associated *E. coli* genomes based on WGA core genome, gene and VF content. **Figure S3.** Manhattan plots of Fisher’s exact test p-values for the OG/VF pathotype (MAEC/commensal isolates) and phylogroup (A/B1) associations. **Figure S4.** Gene organization of the AMR-SSuT/Tn*10* gene cluster. **Figure S5.** Circular genome diagrams for the MAEC 1303 and ECC-1470 replicons with GI and prophage positions. **Figure S6.** Heatmap of VF presence/absence, including gene names/locus tags. **Figure S7.** Gene organization of the Flag-2 and ECC-1470 T6SS/1 gene clusters. (PDF 18 MB)

**Additional file 4: Dataset S1.** This zip archive contains the binary presence/absence matrix of 13,481 OGs in the 25 bovine-associated *E. coli* genomes and the R script for the Fisher’s exact tests to test the associations of OGs/VFs with either pathotype or phylogroup (A/B1). The R script includes also code to create the Manhattan plots in Figure S3. The binary presence/absence matrix of virulence-associated genes needed as second input for the R script is enclosed in Additional file 14: Dataset S7. (ZIP 16 KB)

**Additional file 5: Table S3.** This file includes the SRA study accession numbers for the Illumina and 454 raw reads of the 14 *E. coli* genomes of this study. Additionally it lists the assembly statistics for all 23 bovine-associated *E. coli* draft genomes. (XSLX 8 KB)

**Additional file 6: Dataset S2.** Singleton OGs in the 25 bovine-associated *E. coli* genomes. (XSLX 248 KB)

**Additional file 7: Dataset S3.** This file includes the pathotype-enriched OGs (MAEC or commensal isolates) with a 70% inclusion and 30% exclusion cutoff and their potential association with phylogroup-enriched categories or soft core genomes. OGs significantly associated according to Fisher’s exact test and with a Bonferroni correction are indicated. It also specifies the pathotype group soft core genome and OGs classified as underrepresented and unspecific. For each OG the locus tag and annotation of one representative protein from one *E. coli* genome of the group is shown (or in the case of paralogs several representative proteins). (XSLX 523 KB)

**Additional file 8: Dataset S4.** Phylogroup-enriched OGs (A, B1, B2, or E) with Fisher exact test p-values (and Bonferroni correction) for phylogroup A versus B1 associations and vice versa. Furthermore the file includes the phylogroup group soft core, underrepresented, and unspecific OGs. (XSLX 541 KB)

**Additional file 9: Table S4.** All-strain soft core genome with 70% inclusion cutoff. (XSLX 185 KB)

**Additional file 10: Dataset S5.** Predicted GIs and prophages of MAEC 1303. (XSLX 68 KB)

**Additional file 11: Dataset S6.** Predicted GIs and prophages of MAEC ECC-1470. (XSLX 46 KB)

**Additional file 12: Table S5.** This file contains the overview of the VF panel. Presence (‘1’) and absence (‘0’) of the virulence-associated genes in the 25 bovine-associated *E. coli* genomes is indicated in column “present_in_strain_panel”. Virulence-associated genes were collected from the Virulence Factors Database (VFDB) or from the primary literature (‘own’ in column “source”). (XSLX 55 KB)

**Additional file 13: Table S6.** BLASTP+ hit results for the VF panel in the 25 bovine-associated *coli* genomes. (XSLX 365 KB)

**Additional file 14: Dataset S7.** This zip archive contains the binary presence/absence matrix of virulence-associated genes in the 25 bovine-associated *E. coli* genomes, the VF content clustering cladogram in NEWICK format, and the R script to create the heatmaps in Figure 4 and S4. (ZIP 6 KB)

**Additional file 15: Table S7.** All-strain soft core VF set with 70% inclusion cutoff. (XSLX 10 KB)

**Additional file 16: Dataset S8.** This file includes virulence-associated genes with significant Fisher’s exact test p-values (and Bonferroni correction), which tested the association of VFs with either pathotype (MAEC or commensal strains). Additionally, the pathotype-enriched virulence-associated genes (MAEC or commensal isolates) with a 70% inclusion and 30% exclusion cutoff and their potential association with phylogroup-enriched categories or soft core genomes are listed. It also specifies the pathotype group soft core VF set and virulence-associated genes classified as underrepresented and unspecific. (XSLX 29 KB)

**Additional file 17: Dataset S9.** Significant virulence associated genes (with and without Bonferroni correction) with phylogroup A versus B1 according to Fisher’s exact test. Moreover, phylogroup-enriched virulence-associated genes (A, B1, B2, or E), phylogroup group soft core VF set, underrepresented, and unspecific virulence-associated genes are specified. (XSLX 46 KB)

**Additional file 18: Dataset S10.** BLASTP+ hit results for the *pga* and *paa* gene regions and binary presence/absence matrix in the 25 bovine-associated *E. coli* genomes. The spreadsheet file includes also Fisher’s exact test p-values for significant associated genes to either pathotype or phylogroup. (XSLX 40 KB)

**Additional file 19: Dataset S11.** Pathotype group soft core and unspecific categorisation of the *fec*, *paa*, and *pga* gene regions in the 13 phylogroup A bovine-associated *E. coli* genomes. (XSLX 8 KB)

**Additional file 20: Dataset S12.** Pathotype-enriched (with Fisher’s exact test p-values), group soft core, and unspecific categorisation of the *fec*, *paa*, and *pga* gene regions in the ten phylogroup B1 bovine-associated *E. coli* genomes. (XSLX 9 KB)

**Additional file 21.** Detailed Material & Methods description. (DOCX 113 KB)

**Additional file 22: Dataset S13.** This zip archive contains the GENBANK files with the reannotations of five of the eleven reference bovine-associated *E. coli* genomes. Included are *E. coli* strains AA86, D6-113.11, D6-117.07, D6-117.29, and ECA-727. (ZIP 15 MB)

**Additional file 23: Dataset S14.** This zip archive contains the GENBANK files with the reannotations of six of the eleven reference bovine-associated *E. coli* genomes. Included are *E. coli* strains ECA-O157, ECC-Z, O32:H37 P4, P4-NR, O157:H43 T22, and W26. (ZIP 18 MB)

### Abbreviations

AIEC: adherent invasive *E. coli;*
AMR-SSuT: antimicrobial multidrug resistance to streptomycin, sulfonamide, and tetracycline;
APEC: avian pathogenic *E. coli*;
BRIG: BLAST Ring Image Generator;
CC: clonal complex;
CDS: coding DNA sequence;
CU: chaperone usher pathway fimbriae;
EAEC: enteroaggregative *E. coli*;
ECOR: *E. coli* Reference;
ECP: *E. coli* common pilus;
EHEC: enterohaemorrhagic *E. coli*;
EPEC: enteropathogenic *E. coli*;
ETEC: enterotoxigenic *E. coli*;
ETT2: *E. coli* type III secretion system 2;
ExPEC: extraintestinal pathogenic *E. coli*;
Fec: ferric iron(III)-dicitrate uptake system;
FF: fitness factor;
Flag-1: *E. coli* peritrichous flagella 1 gene cluster;
Flag-2: *E. coli* lateral flagella 2 gene cluster;
G4C: group 4 capsule;
GI: genomic island;
GTR: generalized time-reversible;
HGT: horizontal gene transfer;
IPEC: intestinal pathogenic *E. coli*;
IS: insertion sequence;
LEE: locus of enterocyte effacement;
LPS: lipopolysaccharide;
MAEC: mastitis-associated *E. coli*;
MGE: mobile genetic element;
ML: maximum likelihood;
MLST: multi-locus sequence typing;
MNEC: newborn meningitis-associated *E. coli*;
MPEC: mammary pathogenic *E. coli*;
MST: minimum spanning tree;
OG: orthologous group;
OMP: outer membrane protein;
ORF: open reading frame;
PAI: pathogenicity island;
PAMP: pathogen-associated molecular pattern;
PE: paired-end;
PHAST: PHAge Search Tool;
PTS: phosphotransferase system;
Rhs: rearrangement hotspot;
SLV: single locus variant;
SPATE: serine protease autotransporters of *Enterobacteriaceae*;
SRA: Sequence Read Archive;
ST: sequence type;
T2SS: type II secretion system;
T3SS: type III secretion system;
T5SS: type V secretion system;
T6SS: type VI secretion system;
UPEC: uropathogenic *E. coli*;
VFDB: Virulence Factors Database;
VF: virulence factor;
WGA: whole genome nucleotide alignment.

## Declarations

**Ethics approval and consent to participate**

Not applicable.

**Consent for publication**

Not applicable.

## Availability of data and materials

All raw reads of this study (454 and Illumina) can be accessed from NCBI’s SRA. The corresponding SRA study accession numbers are listed in Additional File 5: Table S3. The assembled and annotated genomes of this study have been deposited at DDBJ/ENA/GenBank under the accession numbers listed in Additional file 1: Table S1. The reannotated sequence files of the eleven bovine-associated reference *E. coli* are available in Additional file 22: Dataset S13 and Additional file 23: Dataset S14. The *E. coli* VF panel is available in the GitHub repository ecoli_VF_collection, https://github.com/aleimba/ecoli_VF_collection [41].

The R scripts used in this study can be found in Additional file 4: Dataset S1 and Additional file 14: Dataset S7. R packages used are mentioned in the methods chapter. Perl scripts are stored in GitHub repository bac-genomics-scripts, https://github.com/aleimba/bac-genomics-scripts [82]. These are licensed under GNU GPLv3. Many of these depend on the BioPerl (v1.006923) module collection [106]. All other data sets supporting the results of this article are included within the article and its additional files.

## Competing interests

The authors declare that they have no competing interests.

## Funding

This work, including the efforts of AL and UD, was funded by Deutsche Forschungsgemeinschaft (DFG) (DO 789/3-1 and DO 789/4-1).

## Authors’ contributions

Conceptualization: AL UD.

Data curation: AL AP.

Formal analysis: AL JV DG.

Funding acquisition: UD.

Investigation: AL.

Methodology: AL.

Project administration: AL.

Resources: AL RD UD.

Software: AL.

Supervision: AL UD.

Validation: AL.

Visualization: AL.

Writing – original draft: AL.

Writing – review & editing: AL UD JV RD AP DG.

All authors read and approved the final manuscript.

## Acknowledgements

We thank Alan McNally for hosting Andreas Leimbach for a week and showing him some of the comparative genomics ropes. We would also like to thank David E. Kerr, Wolfram Petzl, Ynte Schukken, Nahum Shpigel, Olga Wellnitz, Lothar Wieler, and Holm Zerbe for supplying strains.

